# Species-specific enhancement of enterohemorrhagic *E. Coli* pathogenesis mediated by microbiome metabolites

**DOI:** 10.1101/513614

**Authors:** Alessio Tovaglieri, Alexandra Sontheimer-Phelps, Annelies Geirnaert, Rachelle Prantil-Baun, Diogo M. Camacho, David B. Chou, Sasan Jalili-Firoozinezhad, Tomás de Wouters, Magdalena Kasendra, Michael Super, Mark Cartwright, Camilla A. Richmond, David T. Breault, Christophe Lacroix, Donald E. Ingber

## Abstract

**Background:** Species-specific differences in tolerance to infection are exemplified by the high susceptibility of humans to enterohemorrhagic *E. coli* (EHEC) infection whereas mice are relatively resistant to this pathogen. This intrinsic species-specific difference in EHEC infection limits the translation of murine research to human. Furthermore, studying the mechanisms underlying this differential susceptibility is a difficult problem due to complex *in vivo* interactions between the host, pathogen, and disparate commensal microbial communities.

**Results:** We utilize organ-on-a-chip (Organ Chip) microfluidic culture technology to model damage of the human colonic epithelium induced by EHEC infection, and show that epithelial injury is greater when exposed to metabolites derived from the human gut microbiome compared to mouse. Using a multi-omics approach, we discovered four human microbiome metabolites — 4-methyl benzoic acid, 3,4-dimethylbenzoic acid, hexanoic acid, and heptanoic acid — that are sufficient to mediate this effect. The active human microbiome metabolites preferentially induce expression of flagellin, a bacterial protein associated with motility of EHEC and increased epithelial injury. Thus, the decreased tolerance to infection observed in humans versus other species may be due in part to the presence of compounds produced by the human intestinal microbiome that actively promote bacterial pathogenicity.

**Conclusion:** Organ on chip technology allowed the identification of specific human microbiome metabolites modulating EHEC pathogenesis. These identified metabolites are sufficient to increase susceptibility to EHEC in our human Colon Chip model and they contribute to species-specific tolerance. This work suggests that higher concentrations of these metabolites could be the reason for higher susceptibility to EHEC infection in certain human populations, such as children. Furthermore, this research lays the foundation for therapeutic-modulation of microbe products in order to prevent and treat human bacterial infection.

## BACKGROUND

Host tolerance to microbial infections varies greatly between different species [1, 2]. In an era of increasing bacterial antibiotic resistance understanding of the molecular basis for these differences could lead to development of novel tolerance-inducing therapeutic approaches to treat pathogenic infections. For example, in the mammalian intestine, up to 100 trillion commensal bacteria influence host health, and the intestinal microbiome differentially modulates sensitivity to infection in different species [3]. An exquisite example is the difference in tolerance to enterohemorrhagic *E. coli* (EHEC), which causes more than 100,000 infections per year in the USA [4], and can result in development of severe bloody diarrhea, hemorrhagic colitis, and hemolytic uremic syndrome (HUS). The infectious dose for EHEC is 100,000-fold higher in mice compared with humans (10^7^ versus 10^2^ microbes), and even then mice need to be depleted of their microbiome to show symptoms of infection [5–7]. This difference may be due in part to differential localization and expression of receptors for pathogen virulence factors, such as shiga toxin produced by EHEC [8–10], though more human-relevant studies are needed because no animal model fully recapitulates human symptoms of EHEC infection [11]. These dissimilarities in EHEC infection limit the translational potential of murine research to human. Furthermore, studies in human patients are very limited and focused on blood and fecal samples content but do not allow for more in-depth mechanistic investigation of EHEC infection within the gastrointestinal tract. There is indeed a strong need for models that more closely mimic human intestinal pathophysiology in context of bacterial infection.

Additional evidence suggests that products of commensal bacteria are important modulators of host pathophysiology. Metabolites generated by the host gut microbiome, such as acetate produced by *Bifidobacteria*, have been shown to confer protection against EHEC infection in mice [12]. Propionate produced by *Bacteroides* directly inhibits pathogen growth *in vitro* by disrupting intracellular pH homeostasis, and chemically increasing intestinal propionate levels protects mice from *S. Typhimurium* [13]. Butyrate, another short-chain fatty acid produced by commensal bacteria, also modulates host intestinal barrier, injury response and immunity, as well as pathogenicity of EHEC, by altering expression of genes involved in virulence and flagellar motility [14, 15]. Nevertheless, only a few tolerance-modulating microbial metabolites have been identified from fecal samples, and their effects have never been demonstrated in human intestine. In fact, most research on human host-microbiome-pathogen interactions relies on correlative genomic or meta-genomic studies, making identification of causality in humans extremely difficult [16]. Studies also have been carried out by repopulating gnotobiotic mice with complex mixtures of living human versus mouse commensal microbes, but it is extremely difficult to identify soluble metabolites that mediate their effects [3, 17]. Thus, there is a great need for a human model of EHEC infection where contributions of the complex gut microbiome, and specifically soluble metabolites produced by these commensal microbes, can be explored experimentally.

In the present study, we confronted this challenge using human organ-on-a-chip (Organ Chip) microfluidic cell culture technology [18], which can be used to recapitulate human physiology and model various human diseases *in vitro* [19–25]. To explore whether species-specific intestinal microbiomes can influence host tolerance to EHEC infection, we developed a two-channel Colon Chip lined by primary human colon epithelial cells isolated from patient-derived organoids interfaced with human intestinal microvascular endothelial cells (HIMECs), using a recently described technique [26]. The human Colon Chips were infected with EHEC in the presence of soluble metabolites isolated from bioreactor cultures of complex populations of murine or human intestinal commensal microbes. These studies recapitulated the enhanced sensitivity of human colonic epithelium to EHEC in the presence of human microbiome products compared to those from mouse. Surprisingly, however, we discovered that the human microbiome metabolites increased EHEC’s ability to induce epithelial damage, rather than the mouse microbiome products protecting against the damaging effects of this infectious pathogen.

## RESULTS

### Microbiome metabolites recapitulate species-specific tolerance in Colon Chips

To explore how gut microbiome metabolites contribute to species-specific differences in response to infection by EHEC, we cultured healthy, primary, human colon epithelial cells isolated from human donor-derived organoids in close apposition to primary HIMECs within a microfluidic culture device to create a human Colon Chip. This device contains two parallel microchannels separated by a porous (7 μm diameter) extracellular matrix (ECM)-coated membrane; the epithelial cells were cultured on the upper surface of the membrane in the top ‘intestinal lumenal’ channel with gut-specific cell culture medium in both channels, as previously described [26]. The HIMECs were seeded on the opposite side of the same membrane in the lower ‘vascular’ channel (**Fig. 1A**) with gut-specific cell culture medium being perfused through the intestinal luminal channel, and the same medium supplemented with endothelial cell supplements and growth factors through the vascular channel (see **Methods** for details). These culture conditions induced formation of a continuous, undulating, colonic epithelium (**Fig. 1A,B**) that extended across the entire surface of the ECM-coated membrane within 1 week of culture, as visualized using immunofluorescence microscopy (**Fig. 1A**, **right**), phase contrast (**Fig. 1B**) or pseudo-colored imaging (**Fig. 1C**). Human microbiome metabolites (Hmm) or mouse microbiome metabolites (Mmm) were collected from PolyFermS continuous intestinal fermentation bioreactors [27–30] in which complex mouse or human microbiome samples were cultured for 2 weeks under conditions that mimic the internal milieu of the colon (Poeker et al., 2018; Poeker et al., submitted) (see **Methods** for details) (**Fig. 1A**). On day 8 of the Colon Chip culture, the luminal culture medium was replaced with the same medium supplemented with human or murine microbiome metabolites (diluted 1:20 in a PBS-water based solution to 300 mOsm kg^-1^), while continuing to flow the same endothelial culture medium through the vascular channel. Perfusion was continued for 24 hours, followed by introduction of EHEC (1.7 x 10^5^) into the apical lumen in the same medium for 3 hours under static conditions to allow for bacterial cell attachment; medium flow was then re-established and continued for 24 additional hours.

**Figure 1.**
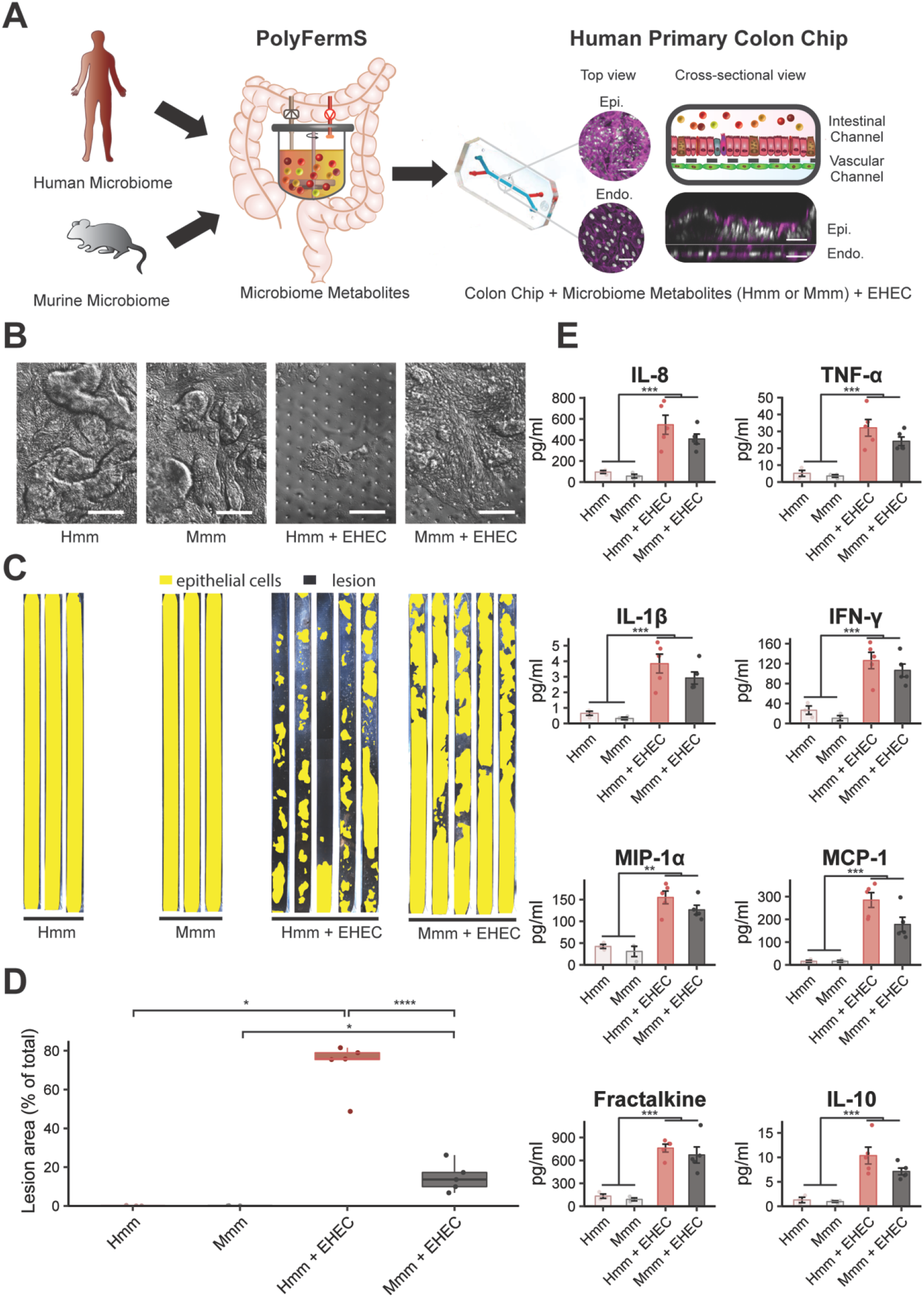
Microbiome metabolites recapitulate species-specific tolerance in Colon Chips. **(A)** A schematic representation of the experimental design illustrating how human or mouse intestinal microbiome metabolites were added to the Intestinal Channel (red) of optically clear, human Colon Chips that are lined by primary human colon epithelial cells (Epi) and directly apposed to a second parallel vascular microchannel (blue) in which HIMVECs (Endo.) are cultured; the two channels are separated by a thin, porous, ECM-coated membrane; F-actin filaments in epithelial and endothelial cells were stained with Phalloidin (magenta) and nuclei with DAPI (white). (**B-D**) Analysis of EHEC-induced epithelial injury on-chip. (**B**) Representative differential interference contrast (DIC) images of the colonic epithelium in the presence of Hmm or Mmm in the presence or absence of EHEC (bar, 100 μm). (**C**) Pseudo-colored images of the entire colon epithelium within the upper channel of the Colon Chip (yellow) cultured in the presence of Hmm or Mmm with or without EHEC (dark regions indicate lesion areas). (**D**) Quantification of epithelial lesion areas under the experimental conditions described in **B** and **C**. (**E**) Changes in levels of various indicated cytokines released into the vascular channel of the Colon Chips by cells cultured under the conditions described in **B** and **C**. *p<0.5, **p<0.01, ***p<0.001, ****p<0.0001.

Using this approach, we were able to recapitulate the increased sensitivity of the human Colon Chip to EHEC infection in the presence of microbiome metabolites from human (Hmm) compared to infection when Mmm were present. This was demonstrated by a greatly increased loss of epithelial cells from the normally continuous colon epithelium in the upper channel of chips exposed to Hmm versus Mmm (**Fig. 1B,C**), which corresponded to over a 5-fold increase in lesion area when quantified using computerized image analysis (**Fig. 1D**). Importantly, control studies confirmed that addition of Mmm or Hmm alone, in the absence of EHEC, did not cause any damage to the colonic epithelium (**Fig. 1B-D**). Thus, intestinal microbiome metabolites are sufficient to recapitulate the species-specific effects on tolerance to EHEC infection observed in previous studies comparing humans and mice, even in the absence of live commensal microbes or immune cells. These findings also indicate that the effects seem to be mediated by direct interactions between the infectious pathogens and the colon epithelium, and not secondary to effects on immune cells that were not present in this study.

### EHEC-induced release of cytokines in the Colon Chip is similar to *in vivo*

As the intestinal epithelium and endothelium contribute to the host innate immune response to bacterial pathogens by secreting inflammatory regulators [31–33], we analyzed the effects of the Hmm and Mmm on secretion of multiple pro-inflammatory cytokines and chemokines, as well as the anti-inflammatory cytokine, interleukin-10 (IL-10), in the presence or absence of EHEC infection (**Fig. 1E**). The pro-inflammatory chemokine interleukin-8 (IL-8) is one of the major cytokines secreted by intestinal epithelial cells [34], and it has been shown to be strongly increased during EHEC infection in humans [35–37]. Additionally, the tumor necrosis factor-α (TNF-α) protein is also increased in human patients who develop EHEC-related HUS [35, 38]. We similarly observed a significant increase in abundance of these two cytokines in the human Colon Chip following EHEC infection when Hmm was present, with IL-8 increasing over 5-fold compared to our control. In addition, the inflammatory mediator interleukin-1 β (IL-1β) has been reported to enhance expression of shiga toxin receptors on endothelial cells, which increases EHEC-related toxicity [39], and we similarly observed increased release of IL-1β into the endothelium-lined vascular channel of these human Colon Chips exposed to Hmm. EHEC bacteria secrete molecules that modulate downstream interferon-_γ_ (IFN-_γ_) signaling in epithelial cells as well [40], and remarkably, we also detected higher levels of IFN-γ following EHEC infection on chip. Moreover, EHEC infection increases circulating Macrophage Inflammatory Protein-1 α (MIP-1 α) and monocyte chemoattractant protein-1 (MCP-1) levels in children [41] and the infected Colon Chips displayed similar increases in the levels of both of these leukocyte chemoattractants. Finally, we discovered that fractalkine levels increased almost 6-fold following EHEC infection, suggesting that this the CX_3_C chemokine family member also may be involved EHEC pathogenesis.

The anti-inflammatory cytokine IL-10 is of interest because it has been investigated as a potential therapeutic to reduce acute inflammation in inflammatory bowel disease [42–44]. Perhaps because of the body’s attempt to counter the effects of EHEC infection, IL-10 concentrations in blood increase with the severity of this disease, and very high levels of IL-10 levels appear to correlate with HUS onset [45]. Similarly, in the infected Colon Chips, IL-10 levels increased almost 10-fold and reached levels (~10 pg ml^-1^) comparable to those observed in human patients with severe colitis induced by EHEC [36] (**Fig. 1E**).

Taken together, these results show that the human Colon Chip recapitulates the pro- and anti-inflammatory cytokine profiles induced by EHEC infection when cultured in the presence of metabolites secreted by the human intestinal microbiome (Hmm). Importantly, when we carried out similar studies in the presence of mouse microbiome products, we found that they produced similar increases in the production of these cytokines as Hmm (**Fig. 1E**), suggesting that the cytokine response generated by the human intestinal cells does not appear to be responsible for the differences in epithelial injury that are induced by EHEC infection under these culture conditions.

### Species-specific injury effects are not due to changes in EHEC colonization

Commensal bacteria of the gut microbiome have been previously shown to inhibit microbial infections by interfering with bacterial colonization of the intestinal epithelium via nutrient competition, secretion of antimicrobials, and adhesion exclusion [13, 46]. For this reason, we investigated whether Hmm and Mmm may differ in their ability to prevent EHEC adhesion and colonization in the human Colon Chips. Results of fluorescence microscopic imaging and quantification of live adherent and non-adherent bacterial cells show that neither the overall distribution of the pathogenic EHEC bacteria (**Fig. S1A**) nor their adhesion to the epithelium (**Fig. S1B**) was altered by the presence of Hmm or Mmm. The numbers of non-adherent EHEC bacteria present in the medium effluent collected from the epithelial lumen of the Colon Chip were also comparable in cultures containing Hmm and Mmm (**Fig. S1C**). Thus, metabolites produced by commensal microbes of the human and mouse gut microbiome do not appear to significantly alter EHEC colonization of the intestinal niche in the human Colon Chips.

### Human microbiome metabolites stimulate bacterial motility

Bacterial chemotaxis and motility are intimately linked to pathogenesis as these processes enable bacteria to detect and react to nutrients, sense clues from other bacteria or the host, and reach their preferred niche [47, 48]. The majority of the intestinal pathogens possess many chemotaxis genes [49], suggesting that chemotaxis increases pathogen fitness and virulence in the gastrointestinal tract. For this reason, there is increasing interest in the investigation of chemotaxis and motility as novel targets to fight infections and avoid problems related to antibiotic resistance [50]. Likewise, EHEC possesses chemotaxis and motility machineries that are used to sense molecules of bacterial or host origin, which can modulate bacterial movement (e.g., autoinducer-3, epinephrine, norepinephrine). To investigate whether enhanced EHEC chemotaxis or motility could be responsible for the phenotype we observed, we carried out a transcriptomics analysis of EHEC bacteria present in the intestinal lumenal compartment of human Colon Chips exposed to either Hmm or Mmm.

These studies revealed that multiple EHEC genes are differentially regulated in the presence of Hmm compared to Mmm, with the greatest changes being observed in the bacterial chemotaxis pathway (**Fig. 2A,B**). We investigated genes involved in both cell and flagellar motility (see **Methods** for details) due to their key role in directional cell movement and observed that multiple genes involved these processes were upregulated in the presence of Hmm. These include the *tar* and *tap* genes, which encode methyl-accepting chemotaxis protein (MCP) chemoreceptors; *cheR* and *cheB* that encode proteins that control MCP receptors; and *cheA* and *cheW*, which encode histidine kinase and the linker protein that associates with MCP receptors (**Fig. 2C,D**).

**Figure 2.**
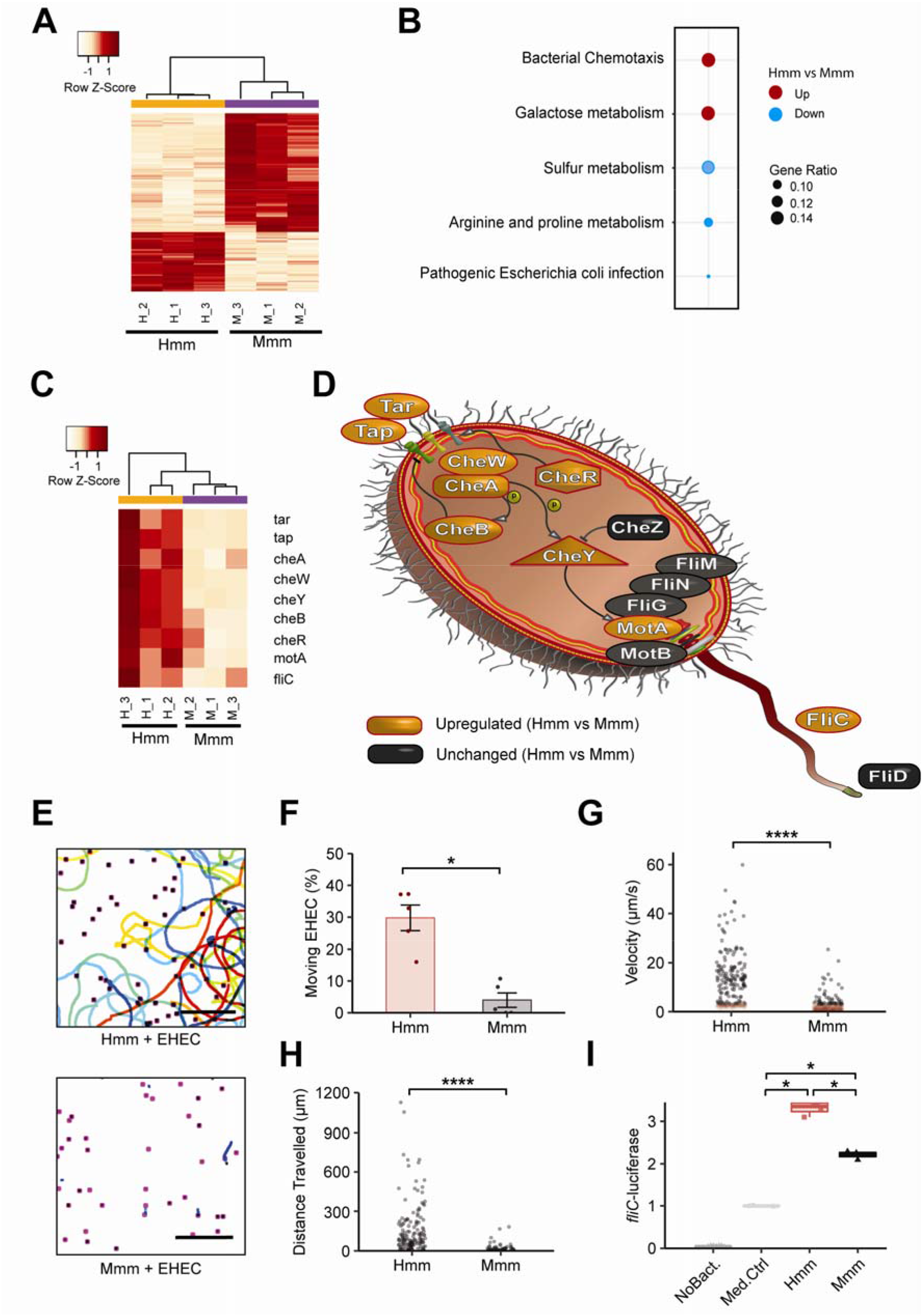
Human microbiome metabolites stimulate bacterial motility. (**A-D**) Changes in the EHEC transcriptome induced by exposure to human (Hmm) versus mouse (Mmm) gut microbiome metabolites. (**A**) Heatmap of differentially expressed genes (red indicates higher levels of expression). (**B**) Gene enrichment analysis. (**C**) Heatmap of chemotaxis and flagellar assembly pathways showing expression levels for relevant motility-related genes in EHEC cultured in the presence of Hmm versus Mmm. (**D**) Schematic of key genes critical in regulating chemotaxis and flagellar assembly in EHEC. (**E**) EHEC swimming motility tracking (lines: bacterial movement tracks; dots: starting points for all tracked bacteria; bar, 100 μm). (**F**) Quantification of the fraction (%) of moving EHEC. (**G**) Mean velocity of each tracked bacterium (red and black: velocity < or > 3 μm s^-1^, respectively). (**H**) Distance traveled (μm) by the moving bacteria. (**I**) *Fli-C*-luciferase expression levels in medium supplemented with Hmm or Mmm [determined by quantifying area under the curve (AUC), and normalizing for the medium control]. *p<0.5, ****p<0.0001.

In addition, the products of *cheA* and *cheW* form a complex that controls a protein encoded by *cheY*, which binds to the flagellar motor and promotes motility when phosphorylated (**Fig. 2D**). All of these genes were upregulated in the presence of Hmm compared to Mmm, and genes that encode components of the flagellar motor system itself, such as *motA* and *fliC* (**Fig. 2D**), were upregulated as well (**Fig. 2C**). Importantly, the transcriptional upregulation of these two latter genes has been directly correlated with higher bacterial motility [51, 52]. Taken together, these results suggest that Hmm preferentially stimulate the motility machinery of EHEC cells relative to Mmm, and that these changes in chemotaxis could mediate the enhanced EHEC-induced pathogenicity we observed in the human Colon Chips.

To further explore this potential mechanism, we evaluated bacterial motility in the presence of Hmm or Mmm using GFP-expressing EHEC bacteria, as previously described [53]. Our results showed a significantly higher fraction of highly motile bacteria in the presence of Hmm (**Fig. 2E,F**) that exhibited both increased velocity and greater distance traveled (**Fig. 2G,H**). Additionally, we also confirmed that expression of the *fliC* transcript was increased by Hmm (**Fig. S2A**) and that the difference in motility was not due to altered EHEC viability (**Fig. S2B**).

To corroborate these findings that Hmm stimulates flagellar motility in EHEC, we screened for metabolites that affect flagellar motility using an EHEC strain expressing a *fliC*-luciferase reporter. Although both Hmm and Mmm increased *fliC*-luciferase expression when added to EHEC, its expression was significantly higher when the human metabolites were present (**Fig. 2I**). There were no distinguishable differences in EHEC *fliC*-luciferase growth under these same conditions, confirming that the detected difference in luciferase signal is not due to differences in cell number (**Fig. S2C**). These results indicate that the presence of metabolites from the human gut microbiome can directly increase EHEC movement by stimulating flagellar motility.

### Identification of specific metabolites that mediate increased pathogenicity

We next used metabolomics analysis to identify specific metabolites in Hmm that are responsible for the effects on EHEC motility we observed. To do this, we compared metabolite levels in the pre-fermentation medium used to culture the microbiome (and that mimics dietary food intake) with the Hmm and Mmm isolated from the final stage of PolyFermS fermentation, which emulate the contents of human proximal colon and mouse cecum colonized by their respective microbiomes. To identify relevant metabolites, we mined our metabolomics data set for metabolites produced by commensal bacteria (*i.e.*, metabolites whose levels increase during the fermentation process), and within these we identified the ones enriched in either the Hmm or Mmm samples. This resulted in identification of a total of 426 metabolites produced by commensal bacteria that were differentially expressed between Hmm and Mmm (**Fig. 3A,B**).

**Figure 3.**
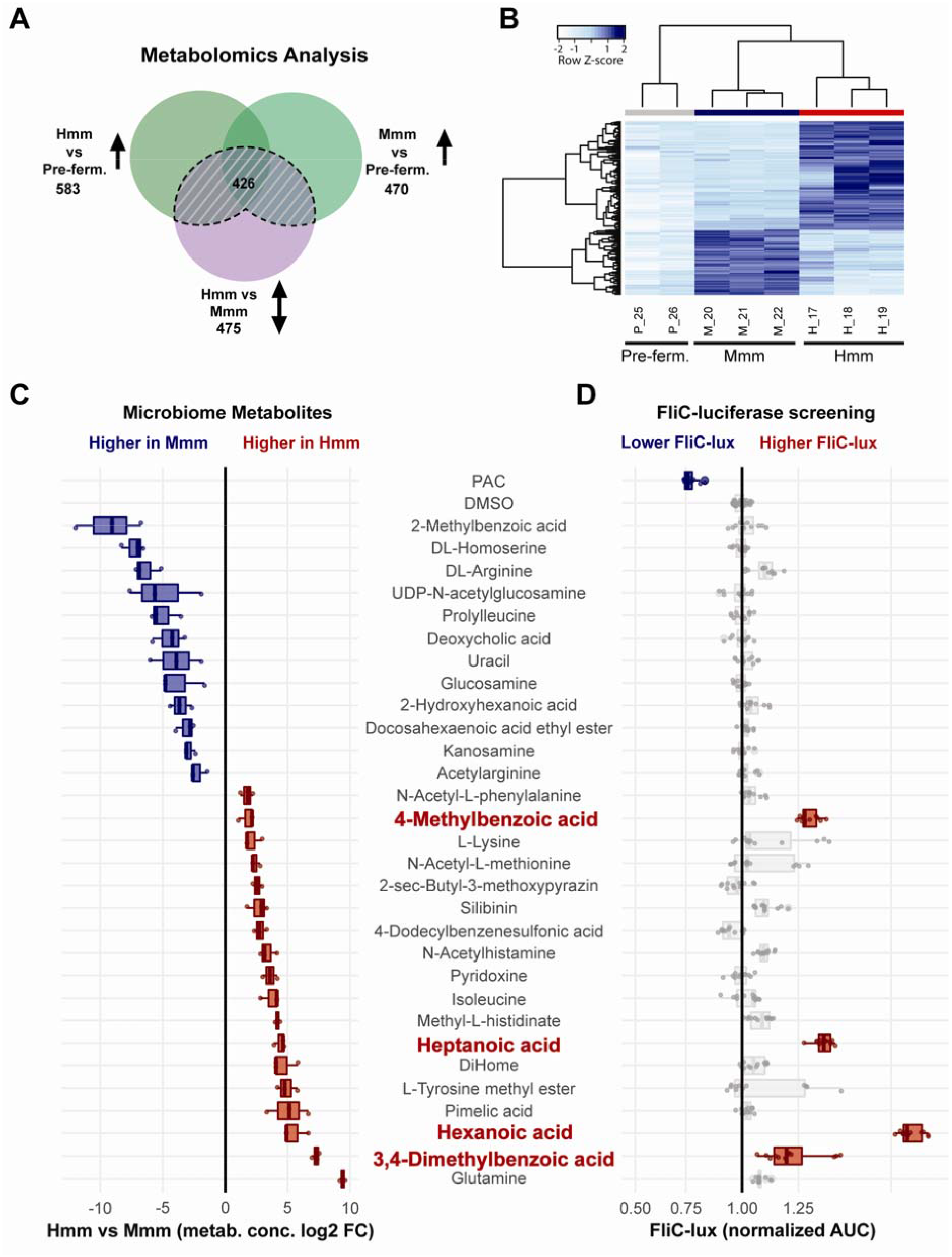
Identification of specific metabolites that mediate EHEC motility. **(A-C)**Results of metabolomics analysis of human versus mouse gut microbiome metabolites. (**A**) Venn-diagram illustrating metabolomics analysis workflow and total numbers of compounds identified in the Hmm and Mmm samples compared to the pre-fermentation medium (Pre-ferm.). (**B**) Heatmap of 426 compounds produced by commensal bacteria that were differentially abundant in human (Hmm) versus mouse (Mmm) microbiome metabolites. (**C**) Relative abundance of 30 microbiome metabolites that were tested (blue and red: higher levels in Mmm or Hmm, respectively). (**D**) Results of *FliC-* luciferase (*FliC*-lux) screening for the 30 selected metabolites (*FliC-lux* levels are presented based on quantification of the AUC; grape seed oligomeric proanthocyanidins (PAC) was used as a negative control; the 4 active metabolites that induced higher *FliC* levels are highlighted in red; all values were normalized against the DMSO control).

To assess their influence on bacterial motility, we selected metabolites that were abundant in either Mmm or Hmm based on our metabolomics analysis. From this MSMS (tandem mass spectrometry) analysis, we included all known compounds (95% confidence), and we also included identifiers that were assigned based on the closest MSMS spectrum in the reference database to the analyte (likely a substructure of the original metabolite); however, we excluded synthetic prescription drugs and known antimicrobial compounds. We performed this selection in order to screen for microbial-derived compounds potentially affecting EHEC motility and not merely bacterial proliferation or viability (see **Methods** for details).

From the final set of 30 compounds, we observed that 12 were present at higher levels in Mmm samples while there was a greater abundance of the remaining 18 in the Hmm set (**Fig. 3C**, **Additional Table 1**). To assess the impact of each one of these compounds on flagellar function and bacterial motility, we carried out a *fliC*-luciferase screening assay as previously described [52]. These studies revealed that only 4 of the selected compounds, which were all present at higher levels in the Hmm samples, significantly increased EHEC *fliC*-luciferase expression: 4-methyl benzoic acid, 3,4-dimethylbenzoic, hexanoic acid and heptanoic acid (**Fig. 3D**). In contrast, we did not identify any metabolites in either the Hmm or Mmm samples that reduced *fliC*-luciferase production, even though we could detect inhibition using the known *fliC* inhibitor, proanthocyanidin (PAC) [54] as a positive control (**Fig. 3D**). When we analyzed the effects of each of these 4 compounds individually over a range of concentrations (1 to 250 μM), we again observed consistent *fliC*-luciferase upregulation (**Fig. S3**) and all of these compounds stimulated EHEC motility when tested at 200 μM in a plate-based bacteria motility assay (**Fig. S4**). Importantly, when all 4 of these compounds were combined at this same concentration and added to Colon Chips cultured in the presence of both EHEC and Mmm, we were able to reconstitute the higher levels of epithelial injury we previously observed in the presence of the complex mixture of Hmm (**Fig. 4A-C**). Taken together, these results show that these four metabolites produced by the human gut microbiome are responsible for increasing EHEC pathogenicity and enhancing epithelial injury in this model.

**Figure 4.**
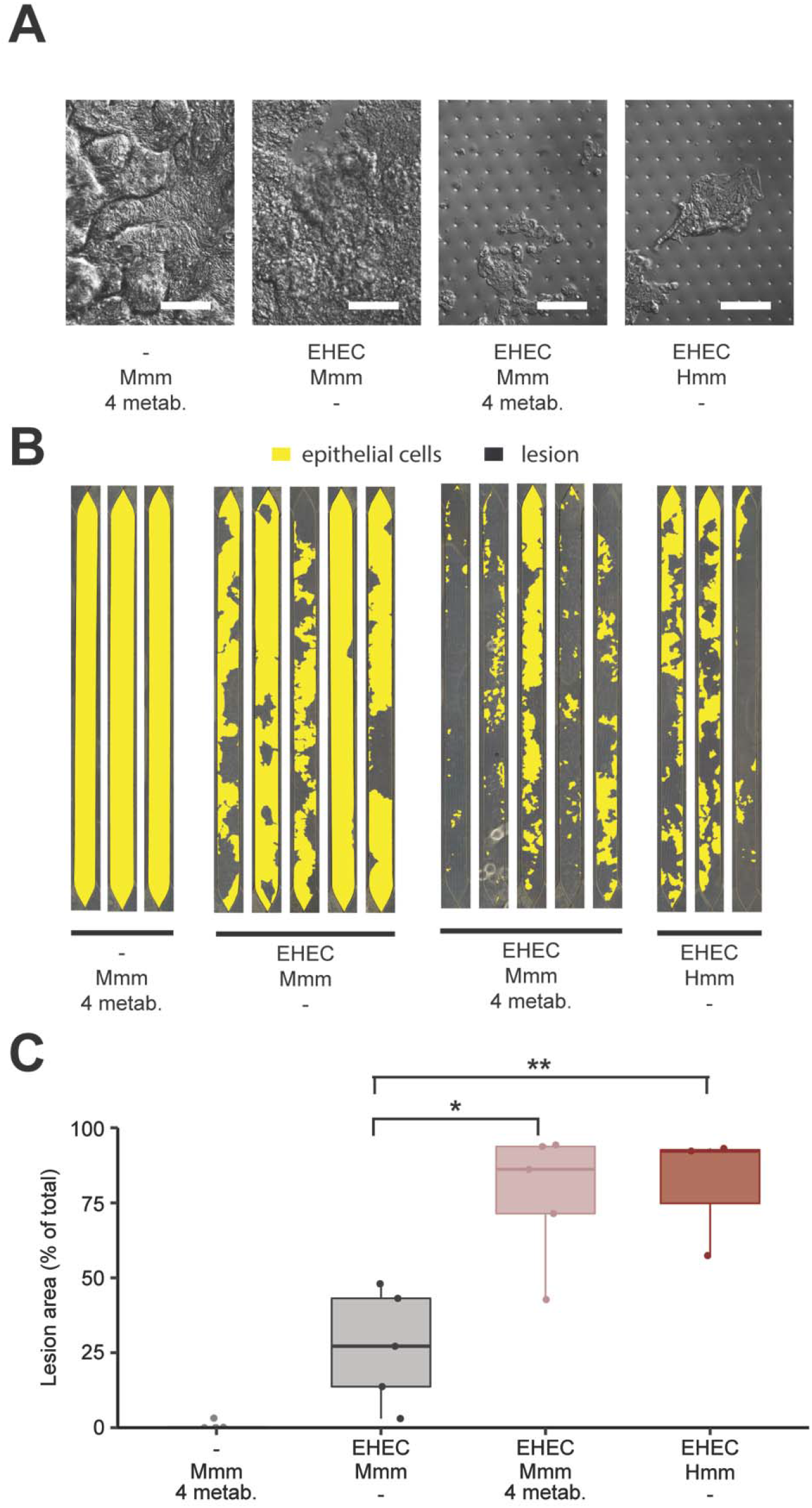
The identified active metabolites mediate increased pathogenicity. (**A-C**) Effect of 3,4-dimethylbenzoic acid, 4-methylbenzoic acid, hexanoic acid, heptanoic acid (4 metab.) on epithelial injury in the Colon Chip in the presence or absence of EHEC, with or without Mmm, compared to the effects of Hmm with EHEC. (**A**) Representative DIC images of the colon epithelium under the various experimental conditions (bar, 100 μm). (**B**) Pseudocolored view of the entire epithelial layer in the Colon Chip (yellow) under the same conditions. (**C**) Quantification of epithelial lesion area size under conditions shown in **B**. *p<0.5, **p<0.01.

## DISCUSSION

In this study, we successfully used Organ Chip technology to model injury of human colon epithelium by infection with EHEC *in vitro*, and to recapitulate species-specific differences in sensitivity to this pathogen by adding either human or mouse gut microbiome metabolites, with greater injury being observed when human metabolites were present. This species-specific effect on epithelial damage occurred despite similar inflammatory signatures in the Colon Chips and was, instead, associated with an increase in expression of EHEC genes associated with known virulence pathways related to chemotaxis and motility. Furthermore, using a metabolomics approach, we were able to identify 4 specific human gut microbiome products that were able to reconstitute the enhanced epithelial injury response observed with complex Hmm; addition of these 4 metabolites also was sufficient to convert the tolerant murine microbiome phenotype into an injury response that mimicked that produced by addition of the human microbiome products. Taken together, these data enhance our understanding of how human commensal bacterial populations modulate intestinal pathogenesis in response to infection, and surprisingly reveal that sometimes this can be a detriment to the host.

Humans are susceptible to EHEC infection at a very low dose (10^2^) [55] whereas the dose required to induce infections in mice is 100,000-fold higher. Moreover, mice need microbiome depletion to show symptoms of infection, further suggesting a potential protective role played by commensal bacteria in murine models [5–7]. Here, using microfluidic Organ Chip technology, we were able to recapitulate this species-specific difference in response to infection observed in humans versus mice *in vivo.* However, we surprisingly discovered that Hmm worsens the outcome of EHEC infection, rather than Mmm inducing a tolerance response. Furthermore, by combining the Organ Chip technology with multi-omics data analysis, we were able to identify specific human microbiome metabolites that exacerbate EHEC-induced injury of the intestinal epithelium independently of effects on bacterial growth or cytokine response. Specifically, we showed that exposure of EHEC to species-specific commensal metabolites changes its chemotaxis-related transcriptional profile and motility behavior, and that the specific four human metabolites we identified upregulate a key motility-related virulence pathway involving flagellar motility. These findings are consistent with those of a past study which showed that flagellin is a key regulator of human intestinal inflammation after EHEC infection *in vivo* [10].

Although we used metabolic analysis to pursue the mechanism by which Hmm and Mmm produce different effects on EHEC-induced epithelial injury, we only focused on known metabolites because these compounds could be obtained commercially and tested experimentally to validate their effects. It is possible that other unknown microbiome-derived metabolites present in the Hmm sample may have additional modulating activities, which could be explored in the future using fractionation of the Hmm sample and in-depth mass spectrometry analysis. Nevertheless, our untargeted metabolomics approach led to the identification of important EHEC injury-enhancing functions for 4 known molecules that were found at significantly higher levels within Hmm versus Mmm. Importantly, a similar experimental approach could be used to identify microbiome-derived modulators of other enteropathogens that exhibit species-specific differences in pathogenicity in the future. This general strategy of comparing how species-specific microbiome metabolites influence host-pathogen interactions in controlled experimental settings could prove to be effective in uncovering new functions for other microbiome metabolites. It also might offer new mechanistic insights into why certain species are more tolerant to specific infectious pathogens than others in the future.

Microbiome metabolites are products of the breakdown of dietary foods and nutrients derived from degradation of host molecules (e.g., mucin) by commensal microbes. Two of the compounds identified (4-methylbenzoic acid, 3,4-dimethylbenzoic acid) are structurally simple phenolic compounds, which can be generated by the degradation of more complex phenolic compounds, such as caffeic acid, ferulic acid and tannins contained in coffee, cereal and grapes [56–58]. Interestingly, these molecules are often promoted as having health benefits based on their anti-oxidant effects; however, our results raise the possibility that they could have negative effects on health if they are metabolized into products that enhance the pathogenicity of EHEC. Taken together, these findings emphasize how further investigation of dietary compounds altering microbiome metabolite production could improve our understanding of how diet alters the intestinal environment and influences patient responses to infection (e.g., by testing the effect of dietary nutrients on the composition and metabolic activity of cultured gut microbiota; Lacroix et al., 2015). The other two compounds we identified that increased epithelial injury are medium chain fatty acids (heptanoic and hexanoic acid). Interestingly, fecal hexanoic acid concentrations have been found to extend over a much higher range in children than in adults [60–62], and it’s known that children are more susceptible to EHEC infection. Taken together with literature showing that short chain fatty acids influence EHEC virulence factor expression and change EHEC infection outcomes in mice [12], these data highlight a key role for microbiome-derived fatty acids in EHEC pathogenesis. Improving our understanding of these host-microbiome-pathogen interactions could provide insight into why certain human groups control EHEC infection while others do not.

This work with Organ Chip technology represents an initial step towards developing advanced *in vitro* human disease models that can enable direct investigation into the cellular and molecular basis of human host-microbiome-pathogen relationships. The human Colon Chip model of EHEC infection described here only included living epithelium and endothelium with soluble microbiome-derived metabolites, and thus, we were not able to examine mechanisms related to direct interactions between living commensal microbes and bacterial pathogens, between the microbes and the host cells, or to address questions centered on contributions of immune cells. However, these are feasible future directions for study as Organ Chip technology can incorporate multiple layers of complexity at the cell, tissue, and organ levels (Ingber, 2016), including co-culture of human intestinal epithelium with complex living gut microbiome [64]. Given that we created these Organ Chips with epithelial cells isolated from patient-derived organoids, this approach also can be used to explore interactions between the host epithelium and commensal microbiome isolated from the same patient, and thus, significantly advance personalized medicine in the future. In addition, this *in vitro* human infection model could be used to develop new biomarkers for susceptibility to infection as well as therapeutics or prophylactics that target microbiome-pathogen interactions, and thereby protect against enteric bacterial infections in an antibiotic-independent manner.

## CONCLUSION

Application of Organ Chip technology to create a human Colon Chip, combined with microbial metabolites isolated from microbial fermentation reactors(PolyFermS), allowed analysis of pathogen-microbiome -human host interactions under in vitro conditions that emulate human intestinal physiology in vivo. The gut microbiome products identified as active were sufficient to recapitulate species-specific differences in response to EHEC infection, with greater injury being observed when human metabolites were present compared to murine metabolites. The four human gut microbiome products we identified – 4-methyl benzoic acid, 3,4-dimethylbenzoic, hexanoic acid, and heptanoic acid – were sufficient to convert the tolerant murine microbiome phenotype into a higher injury response comparable to the one observed in the presence of human microbiome products. This study offers new mechanistic insights in EHEC pathogenesis mediated by human microbiome metabolites, which also could potentially explain different susceptibilities within different human populations, such as the greater sensitivity observed in children. These findings also provide a basis to explore therapeutic and prophylactic modulation of human intestinal contents in order to protect from infection by this potentially life-threatening pathogen.

## METHODS

### EXPERIMENTAL MODEL AND SUBJECT DETAILS

#### PolyFermS

The gut microbial metabolite suspensions used in this study were generated by the PolyFermS platform, a validated continuous in vitro intestinal fermentation model [27](ETH Zürich). Two distinct sources of human and mouse gut microbiome (healthy human adult feces and healthy WT C57BL/6 murine cecal content) were immobilized as described (Poeker et al., 2018, Poeker et al., submitted) and used to inoculate the PolyFermS platform. The PolyFermS pre-fermentation medium for the proximal colon was based on the composition described by Macfarlane et al.,1998 for simulation of the intestinal content (Macfarlane et al. 1998). It includes (g L-1 of distilled water): pectin (citrus) (2), mucine (4), L-cysteine HCl (0.8), bile salts (0.4), KH2PO4 (0.5), NaHCO3 (1.5), NaCl (4.5), KCl (4.5), MgSO4 anhydrated (0.61), CaCl2*2 H2O (0.1), MnCl2* 4 H2O (0.2), FeSO4* 7H20 (0.005), hemin (0.05) and Tween 80 (1 mL). One mL of a filter-sterilized (0.2 μm pore-size) vitamin solution was added to the sterilized (20 min, 120°C) and cooled down medium. The final concentration of vitamins in PolyFermS medium (μg per L) are: Pyridoxine-HCl (Vit. B6) (100); 4-Aminobenzoic acid (PABA) (50); Nicotinic acid (Vit. B3) (50); Biotine (20); Folic acid (20); cyanocobalamin (5); Thiamine (50); riboflavin (50); Phylloquinone (0,075); menadione (10); pantothenate (100). Human pre-fermentation medium included (g L-1 of distilled water): xylan (oat spelts) (2), arabinogalactan (larch) (2), guar gum (1), inulin (1), soluble potato starch (5), casein acid hydrolysate (3), peptone water (5), Bacto tryptone (5), yeast extract (4.5). The PolyFermS pre-fermentation medium for the mouse cecum model was also based on the composition described by Macfarlane et al. (1998), modified for its carbohydrate and protein concentrations for mimicking the chyme of a mouse, accounting for the mouse chow and different digestion physiology (Poeker et al. submitted). Murine prefermentation medium included (g L-1 of distilled water): xylan (oat spelts)(4.8), arabinogalactan (larch) (4.8), soluble corn starch (4), mucine (4), casein acid hydrolysate (3.6), peptone water (6), tryptone (6), yeast extract (5.4) and guar gum was omitted (see **Additional Table 2** for material details).

Human PolyFermS reactors were operated at proximal colon conditions (pH 5.8, 37°C) and a hydraulic retention time of 8 hours (Poeker et al. 2018). Murine PolyFermS reactors were operated at murine cecal conditions (pH 6.5, 37°C) and hydraulic retention time of 12 hours (Poeker at al., submitted). In both models, reactor headspace was continuously flushed with sterile CO2 to control for anaerobiosis. After operating in continuous mode for at least 10 days, microbial composition in both systems remained stable for collection of effluents for the Colon Chip experiments, as indicated by stable base consumption, fermentation metabolite production and bacterial population composition [30]. Fresh effluents were collected from PolyFermS reactor, centrifuged at 16,000 x g for 10 minutes in a pre-cooled centrifuge (Heraeus Biofuge Primo 230, MultiTemp Scientific AG, Kloten, Switzerland) at 4°C. The supernatant was filter sterilized using a 0.2 μm filter (Minisart-plus 0.2 μm filter, Satorius Stedim Biotech GmbH, Goettingen, Germany) and stored in 1 mL aliquots at -80°C until usage.

#### Organoid cultures

Human intestinal epithelium was isolated from resections or endoscopic tissue biopsies from 2 females and one male. Resections, consisting of full thickness pieces of human colon, were obtained anonymously from healthy regions of colonic resection specimens processed in the Department of Pathology at Massachusetts General Hospital under an existing Institutional Review Board approved protocol (#2015P001859). Specimens were restricted to healthy (non-neoplastic) disease samples, and tissue was taken from healthy normal regions as determined by careful gross examination. Endoscopic biopsies were collected from macroscopically normal (grossly unaffected) areas of the colon of de-identified patients undergoing endoscopy for abdominal complaints. Informed consent and developmentally-appropriate assent were obtained at Boston Children’s Hospital from the donors’ guardian and the donor, respectively. All methods were carried out in accordance with the Institutional Review Board of Boston Children’s Hospital (Protocol number IRB-P00000529) approval.

Tissues from the resections were dissected to detach the epithelium, and the epithelial layer or the entire biopsy was digested with 2□mg□ml^−1^ collagenase I for 40□min at 37L°C followed by mechanical dissociation, as previously described [26]. Organoids were grown embedded in expansion medium in growth factor-reduced Matrigel [26, 65]. Expansion medium consists of advanced DMEM F12 supplemented with: L-WRN (Wnt3a, R-spondin, noggin) conditioned medium (65% vol vol^-1^), glutamax, 10mM HEPES, murine epidermal growth factor (50 ng ml^-1^), N2 supplement, B27 supplement, 10 nM human [Leu15]-gastrin I, 1 mM n-acetyl cysteine, 10 mM nicotinamide, 10 μM SB202190, 500 nM A83-01, primocin (100 μg ml^-1^) [65, 66] (see **Additional Table 2** for material details).

#### Endothelial cell culture

Human intestinal microvascular endothelial cells (HIMECs) were obtained from ScienCell (Cat#2900) expanded in Microvascular Endothelial Cell Growth Medium-2 BulletKit (EGM-2MV). Endothelial cells were subcultured less than 5 times before use.

#### Colon Chip Cultures

Organ Chips composed of poly-dimethylsiloxane (PDMS) were obtained from Emulate Inc. (Boston, MA). The Organ Chips consist of two parallel microchannels (1000 x 1000 μm and 1000 x 200 μm; Width x Height) separated by a thin (50 μm) porous membrane (7 μm pore diameter, 40 μm spacing). Organ Chips were provided by Emulate Inc. (Cat#10231-2). The chips were activated using ER-1 and ER-2 solutions from Emulate Inc. (Boston, MA) and then coated with type I collagen (200 μg ml^−1^) and Matrigel (1% in PBS), as described [26]. Colonic organoids were fragmented by incubating for 2 minutes at 37°C in Triple E express diluted in PBS 1:1 (vol:vol) supplemented with 10 μM Y-27632, and seeded on the ECM-coated membrane in the intestinal luminal channel of the Colon Chip (210,000 cells/chip) in expansion medium supplemented with 10 μM Y-27632. Chips were incubated overnight at 37°C in expansion medium. The following day expansion medium was perfused with cell culture medium at 60 μl h^−1^ through top and bottom channels. Medium flow was driven at a constant flow rate using a peristaltic pump (Ismatec; Cat#ISM938D)

After 7 days, HIMEC were seeded in the vascular side of the Colon Chip (250,000 cells/chip) in expansion medium supplemented with human epidermal growth factor, vascular endothelial growth factor, human fibroblastic growth factor-B, R3-Insulin-like growth factor-1, ascorbic acid and no antibiotics. To allow endothelial cells to adhere to the membrane, chips were inverted under static conditions for 1 hour, and then the chips were placed upright and perfusion was restored. After 24 hours, the vascular channel medium was switched to recombinant organoid expansion medium. Meanwhile the intestinal luminal channel medium was switched to 5% (vol vol^-1^) human (Hmm) or mouse (Mmm) gut microbiome metabolites isolated from PolyFermS bioreactors, diluted in phosphate-buffered saline (PBS) containing calcium and magnesium (final osmolarity = 300 mOsm kg^-1^), filtered through a 0.2 μm filter (Corning) and stored at -80°C. Recombinant organoid expansion medium consists of organoid expansion medium except that of L-WRN conditioned medium was substituted with recombinant Wnt-3a (100 ng ml^-1^), murine noggin (100 ng ml^-1^), murine R-spondin-1 (1 μg ml^-1^), and supplemented with human epidermal growth factor, vascular endothelial growth factor, human fibroblastic growth factor-B, R3-Insulin-like growth factor-1, ascorbic acid and no antibiotics(see **Additional Table 2** for material details).

### Bacterial growth conditions

EHEC-GFP was generated from NR-3 *Escherichia coli*, EDL931, serotype O157:H7, transformed with pGEN-GFP(LVA) CbR plasmid. The EHEC *fliC*-luciferase strain was generated by transforming EHEC EDL931 with *fliC*-luciferase plasmid kindly provided by H.L. Mobley, as described [52]. EHEC and EHEC-GFP were grown to 0.5 McFarland (1.5 x 10° CFUs) in RPMI medium supplemented with 10% glucose, centrifuged, and stored at -80°C in saline 10% glycerol, while EHEC *fliC*-luciferase bacteria were grown in Luria Broth (LB) containing (g l^-1^ of distilled water) Bacto Tryptone (10), Bacto Yeast Extract (5) and sodium chloride (10) and the appropriate antibiotic; the following day, the bacteria were diluted or resuspended in medium and at the dose indicated, according to the experimental need (see **Additional Table 2** for material details).

## METHODS DETAILS

### Colon Chip Infection

Colon Chips were cultured in the intestinal lumen channel of the chip in 5% (vol vol^-1^) human or mouse gut microbiome metabolites isolated from PolyFermS bioreactors, diluted in phosphate-buffered saline (PBS; final osmolarity = 300 mOsm kg^-1^) for 24 hours. The following day, the intestinal channel was infected with 1.7 x 10^5^ EHEC-GFP by adding the bacteria to into the channel lumen in medium again with or without Hmm or Mmm. Chips were maintained under static conditions for 3 hours to promote EHEC colonization, and then perfused at 60 μl h^−1^.

### Epithelial lesion analysis

One day post infection, Colon Chips were washed with PBS and fixed with 4% paraformaldehyde in PBS for 2 hours. The Chips were imaged using a Leica DM IL LED microscope and images were stitched together with Basler Phylon Software. The area occupied by cells and the total area of the chip were measured using Fiji software [67].

### *FliC*-luciferase reporter assay

The EHEC bacteria were grown overnight in LB medium diluted 1:1000 and the *fliC*-luciferase assay was carried as described [54]. Compound screening was carried at a concentration of 10 μM of the compound diluted in dimethyl sulfoxide (DMSO; final concentration 0.1% vol vol^-1^) or an equivalent volume of DMSO as a vehicle control. Grape seed oligomeric proanthocyanidins (PAC) was used as negative control at a dose of 100 μg ml^-1^ [54]. All compounds used for motility screening were purchased from MedChemExpress except DiHOME (Cayman Chemicals) and PAC (Sigma). Compounds were dosed from 0.97 μM to 250 μM, followed by sealing the plate with a gas permeable membrane (EK Scientific) and incubating at 37°C in a Synergy HT Microplate Reader. Luciferase luminescence and optical density at 600 nm wavelength (OD600) were measured at 20-minute interval for 11 hours.

### Metabolomics

Samples were centrifuged at 10000g for 5 minutes followed by biphasic chloroform-methanol extraction. All samples were run for untargeted mass spectrometry on a ThermoFisher Q-exactive mass spectrometer (Small Molecule Mass Spectrometry Facility, FAS Division of Science Operations Harvard University). Compound Discovery Software was utilized to assign compound names (95% confidence). If the parent ion was not found, the compound with the closest spectrum was used as an identifier, thus indicating a potential substructure of the original metabolite. In the case of multiple metabolites matching to the same identifier, priority was given to the metabolite identified with the highest average area value. From our analysis, we identified 426 metabolites enriched in either Hmm or Mmm, and selected all the metabolites with an assigned compound name. Within these metabolites, we selected all 30 commercially available compounds, while excluding known synthetic prescription drugs, antimicrobial agents or potential chemical contaminants (**Additional Table 1**) and screened them for their effect on EHEC flagellar motility.

### RNA isolation and gene expression

Endothelial cells were first removed using Trypsin-EDTA (0.25%) from the vascular channel of the Colon Chip. Epithelial cells were then isolated from the intestinal luminal compartment and RNA was isolated using an RNAeasy Mini Kit. For qPCR measurement cDNA was synthetized using SuperScript IV VILO Master Mix (Thermo Fisher Scientific) and primers (**Additional Table 2**) and Powerup SYBR Green Master Mix (Thermo Fisher Scientific) were utilized for amplification. For RNA seq analysis, the RNA concentration was measured using a Qubit instrument and Quant-it reagents (Thermo Fisher Scientific). RNA purity was assessed by measuring the ratio of absorbance at 260/230 nm, and 260/280 nm on a Nanodrop Instrument (Thermo Fisher Scientific). RNA integrity was measured using TapeStation 2200 (Agilent Technologies). Bacterial and human ribosomal RNA was depleted from total RNA samples using an Epidemiology Ribo-Zero Gold rRNA Removal kit (Illumina, Inc.) on an Apollo324 automated workstation (Takara Bio USA). The resulting ribosomal-RNA-depleted RNA samples were immediately converted into stranded Illumina sequencing libraries using 200 bp fragmentation and sequential adapter addition on an Apollo324 automated workstation following manufacturer’s specifications (PrepX RNA-seq for Illumina Library kit, Takara Bio USA). Libraries were enriched and indexed using 15 cycles of amplification (LongAmp Taq 2x MasterMix, New England BioLabs Inc.) with PCR primers which include a 6 bp index sequence to allow for multiplexing (custom oligo order from Integrated DNA Technologies). Excess PCR reagents were removed using magnetic bead-based cleanup on an Apollo324 automated workstation (PCR Clean DX beads, Aline Biosciences). Resulting libraries were assessed using a 2200 TapeStation (Agilent Technologies) and quantified by QPCR (Kapa Biosystems). Libraries were pooled and sequenced on four lanes of a HiSeq 2500 v4 high output flow cell using single end, 50bp reads (Illumina, Inc.).

### Cytokines/chemokines analysis

Levels of cytokines and chemokines within medium collected from the effluent of the vascular channel were measured using MSD U-plex Assay (Meso Scale Diagnostic). Medium samples were collected 6 hours post EHEC infection (3 hours after restoring the flow to the Colon Chip.

### Bacterial Motility Tracking

EHEC-GFP bacteria were grown 6 hours at 37°C in Hmm or Mmm, then transferred to plasma-treated cover slips and imaged using a Zeiss Axio Observer Z1 microscope for 3 minutes, as described [53]. The videos were then processed using Fiji, an image processing package of ImageJ, StackReg to stabilize the video, cropped to remove video edges and particles were tracked using TrackMate plugin [67–69]. Particles tracked for less than 1 second were removed from the analysis. Bacteria with a speed higher then 3 μm s^-1^ were considered motile. TrackMate mean velocity was calculated as the mean of the instantaneous velocity and distance traveled shown over the total video time (3 minutes). For better visualization of the full particle tracks, we changed the color Look-Up Table to have a white background using Fiji and applied a minimum filter in Adobe Photoshop to widen the tracks path and make it visible once the image size was reduced for publication purposes.

### Quantification of bacterial numbers

To quantify adherent bacteria, epithelial cells were washed with PBS and isolated with Trypsin-EDTA (0.25%) – Type IV Collagenase (1 mg ml^-1^) 20 min at 37°C. Adherent bacteria and bacteria contained within samples of medium effluent, collected 6 hours post infection, were diluted and plated on soy agar plates with sheep blood with an Eddy Jet 2 automated spiral plater (UL Instruments). Plates were incubated overnight at 37°C and colony-forming units (CFUs) were quantified using a Flash & Go automatic colony counter (UL Instruments).

### Bacteria viability

EHEC-GFP bacteria were grown 6 hours at 37°C in medium containing Hmm or Mmm, then propidium iodide solution was added at a final concentration of 10 mg ml^-1^ for 5 min at room temperature as reported [70]. Bacterial GFP and propidium iodide were imaged using a Zeiss Axio Observer Z1 microscope; the dead bacteria fraction was calculated as PI positive bacteria divided by the total number of bacteria.

### Bacteria Swimming plate assay

Swimming motility was assessed using 0.25% agar LB plates. Overnight cultures of EHEC-GFP bacteria were standardized at 1 OD600 and 1.5 μl of the culture medium was added to the center of the agar plate with a sterile pipette tip as described [71]. Bacterial swimming was quantified at 12 hours, imaging the plates using a FluorChem M imaging system (ProteinSimple). The area occupied by bacteria was then measured using Fiji [67].

### Colon Chip epithelium and bacteria imaging

Colon Chips infected with EHEC-GFP and uninfected controls were washed with PBS and fixed with 4% paraformaldehyde for 2 hours. Following fixation epithelial cells and bacteria were labelled with Alexa Fluor 647 Phalloidin, 4’,6-Diamidino-2-Phenylindole, Dihydrochloride (DAPI) and anti-green fluorescent protein-Alexa Fluor 488 conjugate. Images were acquired with an inverted laser-scanning confocal microscope (Leica SP5 X MP DMI-6000) and processed using IMARIS.

## QUANTIFICATION AND STATISTICAL ANALYSIS

### Analysis of *fliC*-luciferase reporter assay data

Raw luciferase signal at each time point was divided by OD600 to control for bacterial growth. The area under the curve (AUC) was calculated and all the statistics were performed using R language and environment for statistical computing [72]. Each compound was run in quadruplicate and the screening repeated 3 times. A Mann–Whitney–Wilcoxon test was performed followed by Bonferroni correction for multiple comparisons [73–75]. Significant differences were selected with fold change higher than 20% and adjusted p-value < 0.0001. We then generated dose curves for the newly identified compounds modulating *fliC*-luciferase serially diluting 1:2 from 250 μM to 0.97 μM. For *fliC*-luciferase compound dose curves, each dot indicates the mean and the standard error of the mean (SEM).

### Analysis of metabolomics

Raw data were normalized using R metabolomics package [76] and groups compared using Linear Models for Microarray Data (limma) package [77]. We applied a cut off of 0. 05 on adjusted p-value and fold change greater then 1.5.

### Analysis of RNA-seq data

The data was processed using bcbio-nextgen. We used the STAR alignments, FASTWQ files, and Salmon quantification to generate quality control metrics of the samples [78, 79]. Differential expression was computed using DESeq2 package in R [80]. Genes were considered differentially expressed based on both an absolute fold-change larger than 1.5 and an FDR-corrected p-value cutoff less than 0.05. We performed pathway enrichment on this gene signature using the enrichKEGG function in the clusterProfiler package in R [81]. To compare gene clusters associated with a KEGG pathway, we used the compareCluster function in the clusterProfiler package with a p-value cutoff less than 0.001. Represented pathways are based on the following KEGG IDs from the *E. coli* O157:H7 EDL933 annotations: Bacterial chemotaxis pathway (KEGG ID: ece02030), Cellular motility (KEGG ID: 09142: chemotaxis + flagellar assembly), Pathogenic *E. coli* infection genes (KEGG ID: ece05130), Arginine and proline metabolism (KEGG ID: ece00330), Galactose metabolism (KEGG ID: ece00052), and Sulfur metabolism (KEGG ID: ece00920).

### Statistics

All statistical analysis were carried out in R using custom scripts (see **Additional Table S** for software and algorithms details). The Mann-Whitney test was used to compare lesion area of Colon Chips, cytokines expression, percentage of moving bacteria, their velocity, distance travelled, qPCR and bacterial motility plate assay. For the *fliC-* luciferase screening, significance was calculated using a Mann-Whitney test, with p-values adjusted for multiple comparisons using the Bonferroni method. All boxplots represent median, first and third quartile of the data distribution, with whiskers extending to the largest value no further than 1.5 times the inter-quartile range. Bar plots represent mean value of the data with SEM; dots in bar- or box-plots indicate the sample size *N* for each experiment. For Colon Chips experiments, *N* is equal to the number of chips used; for the *fliC*-luciferase experiment, *N* corresponds to a single well; for the quantification of the fraction of moving bacteria, *N* indicates the number of videos analyzed; for all the other bacterial tracking experiments, *N* indicates a single bacterium tracked; for plate-based swimming assays, *N* indicates an individual plate.

## DECLARATIONS

### Ethics approval and consent to participate

Human colonic resections were obtained anonymously from the Department of Pathology at Massachusetts General Hospital under an existing Institutional Review Board approved protocol (#2015P001859). Endoscopic biopsies were collected from de-identified patients from Boston Children’s Hospital. Informed consent and developmentally-appropriate assent were obtained at Boston Children’s Hospital from the donors’ guardian and the donor, respectively. All methods were carried out in accordance with the Institutional Review Board of Boston Children’s Hospital (Protocol number IRB-P00000529) approval.

### Consent for publication

Not applicable.

### Availability of data and material

RNA-seq data have been deposited to the Sequence Read Archive (accession: PRJNA497914). Further information and requests for resources and reagents should be directed to and will be fulfilled by the Lead Contact, Donald E. Ingber.

## Competing interests

D.E.I. holds equity in Emulate, Inc., consults to the company, and chairs its scientific advisory board; he also is an inventor on relevant patents.

## Funding

This work was funded by Defense Advanced Research Projects Agency under Cooperative Agreement Number W911NF-12-2-0036 (to D.E.I.) and the Wyss Institute for Biologically Inspired Engineering at Harvard University. The commensal bacteria fermentation was supported SINERGIA grant 35150 of the Swiss National Science Foundation (to C.L., A.G and T.d-W). The production of human organoids from was supported by NIH grants R01DK084056 and P30DK034854 (to D.T.B.) and by NIH grant 5T32CA009216-37 (to D.B.C.).

## Autors’ contributions

Conceptualization, A.T., R.P.-B.,M.K., C.L. and D.E.I.; Methodology, A.T., A.S.-P, M.K, D.B.C., A.G., T.d.-W.,D.T.B., C.A.R., C.L. and D.E.I; Software, Data Curation and Formal Analysis, A.T., D.M.C.; Investigation: A.T., A.S.-P., S. J.-F., A.G., T.d.-W.; Resources, D.T.B., C.A.R., D.B.C., M.S, M.C.; Writing – Original Draft, A.T., D.E.I.; Writing – Review & Editing, A.T., D.B.C., D.T.B., C.L., A.G., M.K., R.P.-B., A.S.-P, D.M.C., S. J.-F. and D.E.I; Visualization, A.T., D.B.C and D.E.I., Supervision, R.P.-B, C.L. and D.E.I, Funding Acquisition, D.B.C., D.T.B., C.A.R, M.S., C.L. and D.E.I.

## Acknowledgments

We thank H.L. Mobley for his generous gift of *fliC-lux* plasmid; A. Sanz Garcia and S. Leng for their help with R coding; the Bauer Core at Harvard University, Harvard Chan Bioinformatic Core, C. Vidoudez and S.A. Trauger at the Harvard Small Molecule Mass Spectrometry Core for their support in the data collection; A. Monreal, O. Levy, and A. Chalkiadaki for their expert advice; and T. Ferrante for assistance with imaging.

## SUPPLEMENTARY FIGURE LEGEND

**Figure S1.**
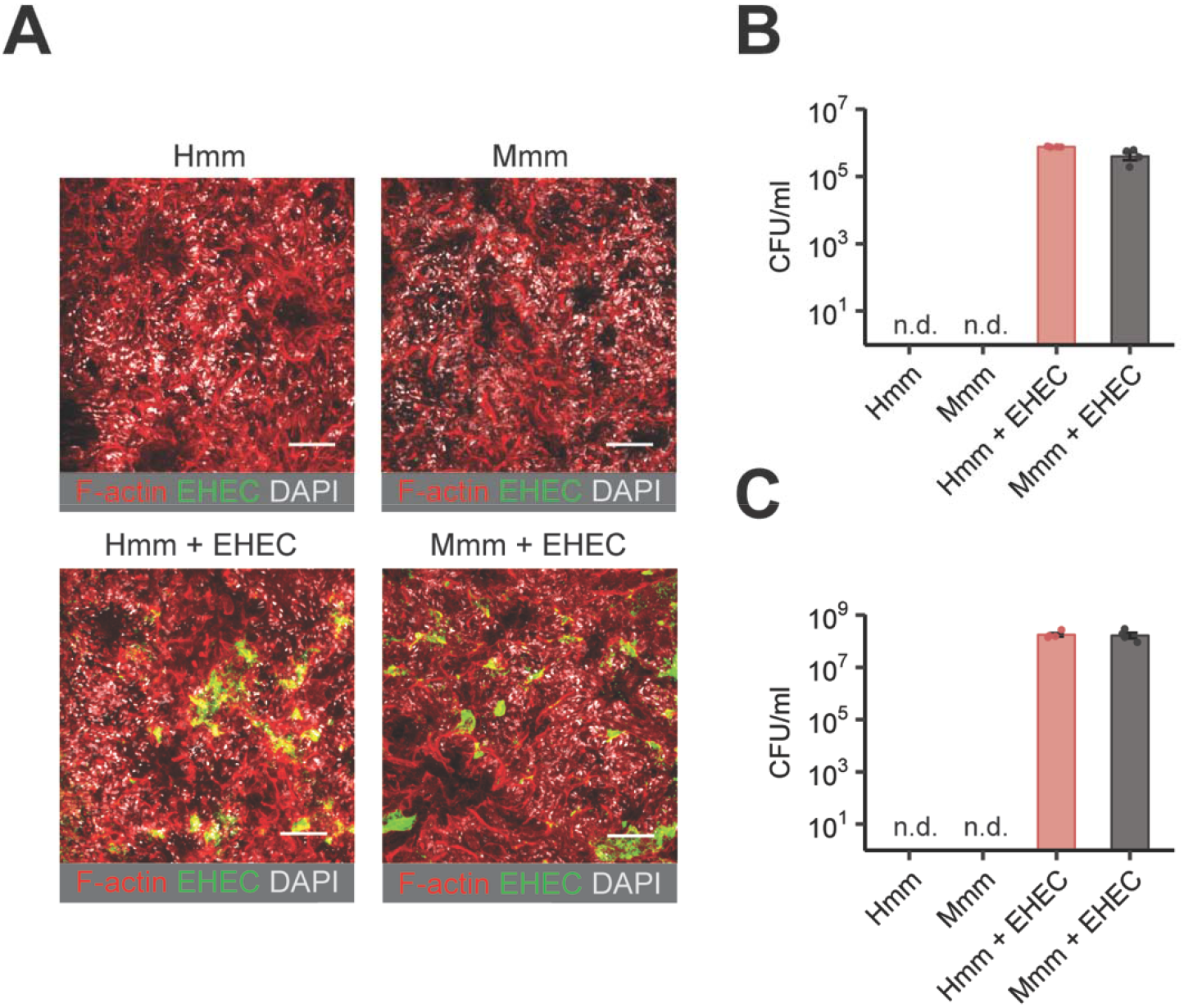
Species-specific injury effects are not due to changes in EHEC colonization. (**A-C**) EHEC colonization of the human Colon Chip. (**A**) Representative fluorescence images showing the epithelial layer of infected and control Colon Chips in the presence or absence of Hmm or Mmm, with or without EHEC present (red: F-actin, green: GFP-EHEC, white: nuclei; bar, 100 μm). (**B**) Quantification of EHEC bacteria adherent to the intestinal epithelium. (**C**) Quantification of non-adherent EHEC quantification floating in the culture medium.

**Figure S2.**
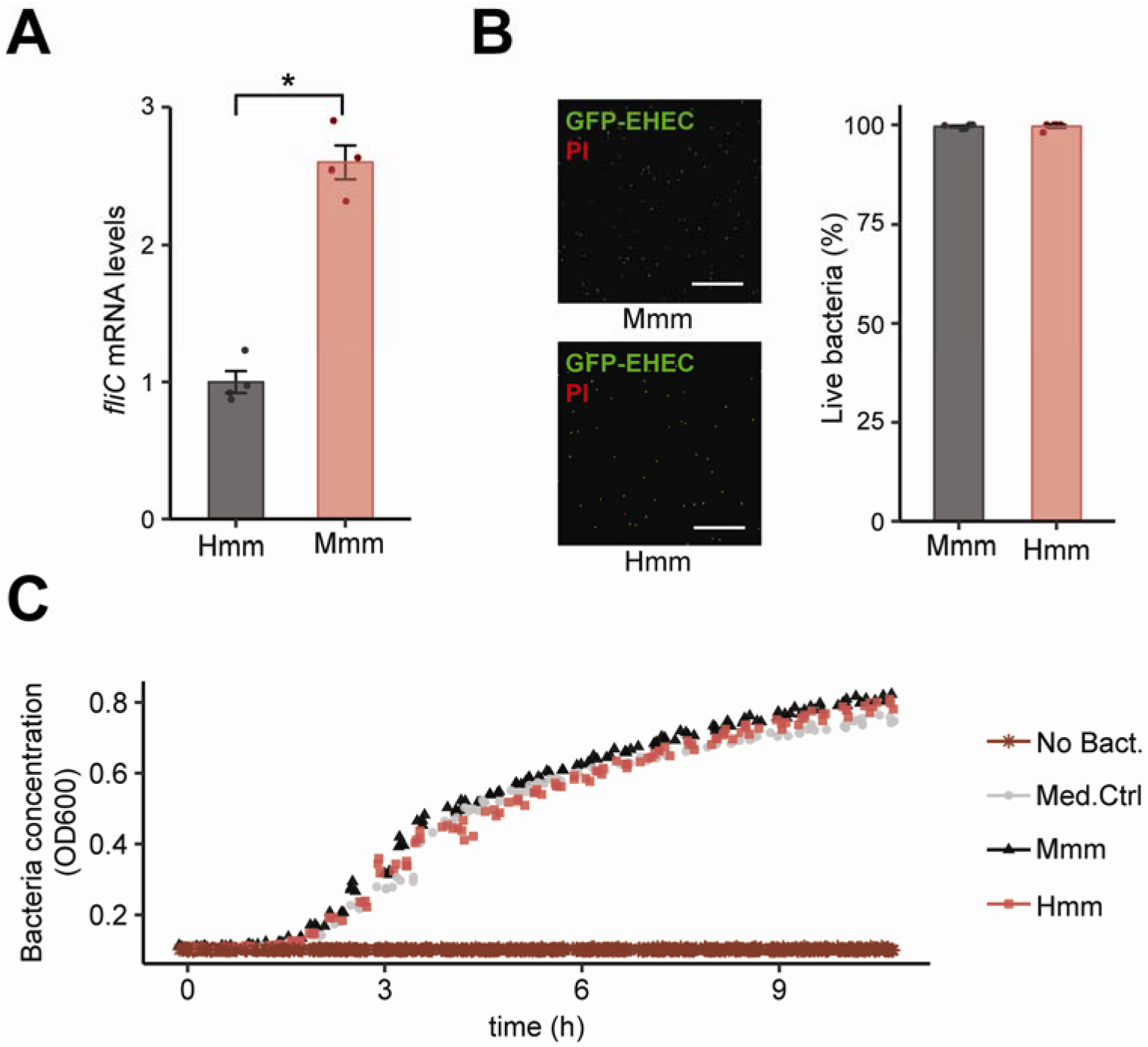
*FliC* gene transcript is upregulated by Hmm, the species-specific motility effect is not due to changes bacteria viability, and the *FliC*-luciferase increase in the presence of Hmm is not due to altered bacterial growth. (**A**) *FliC* mRNA levels in EHEC cultured with Hmm or Mmm (shown as linearized, normalized fold change). (**B**) Fluorescence microscopic image of GFP-EHEC (green) and quantification of EHEC viability by staining with propidium iodide (red; bar, 100 μm). (**C**) Bacterial concentration determined as optical density measured at 600 nm (OD600) of EHEC *fliC*-luciferase in the presence of Mmm or Mmm.

**Figure S3.**
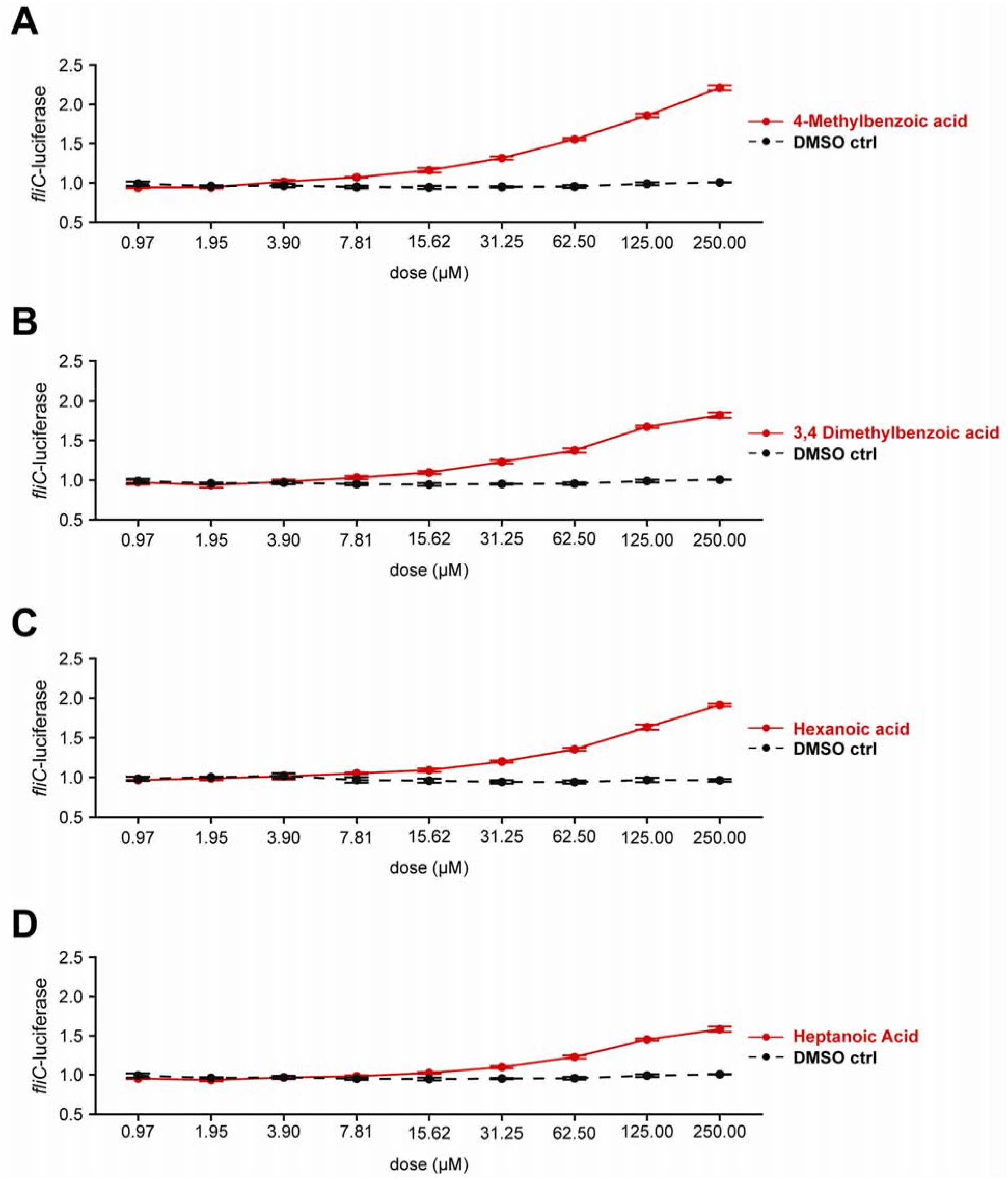
Each of the 4 identified metabolites increases FliC expression in a dose-dependent manner. FliC-luciferase levels (determined by quantifying the AUC and normalizing for the DMSO control) of 4-methylbenzoic acid, 3,4 dimethylbenzoic acid, hexanoic acid, and heptanoic acid metabolites measured at indicated concentrations.

**Figure S4.**
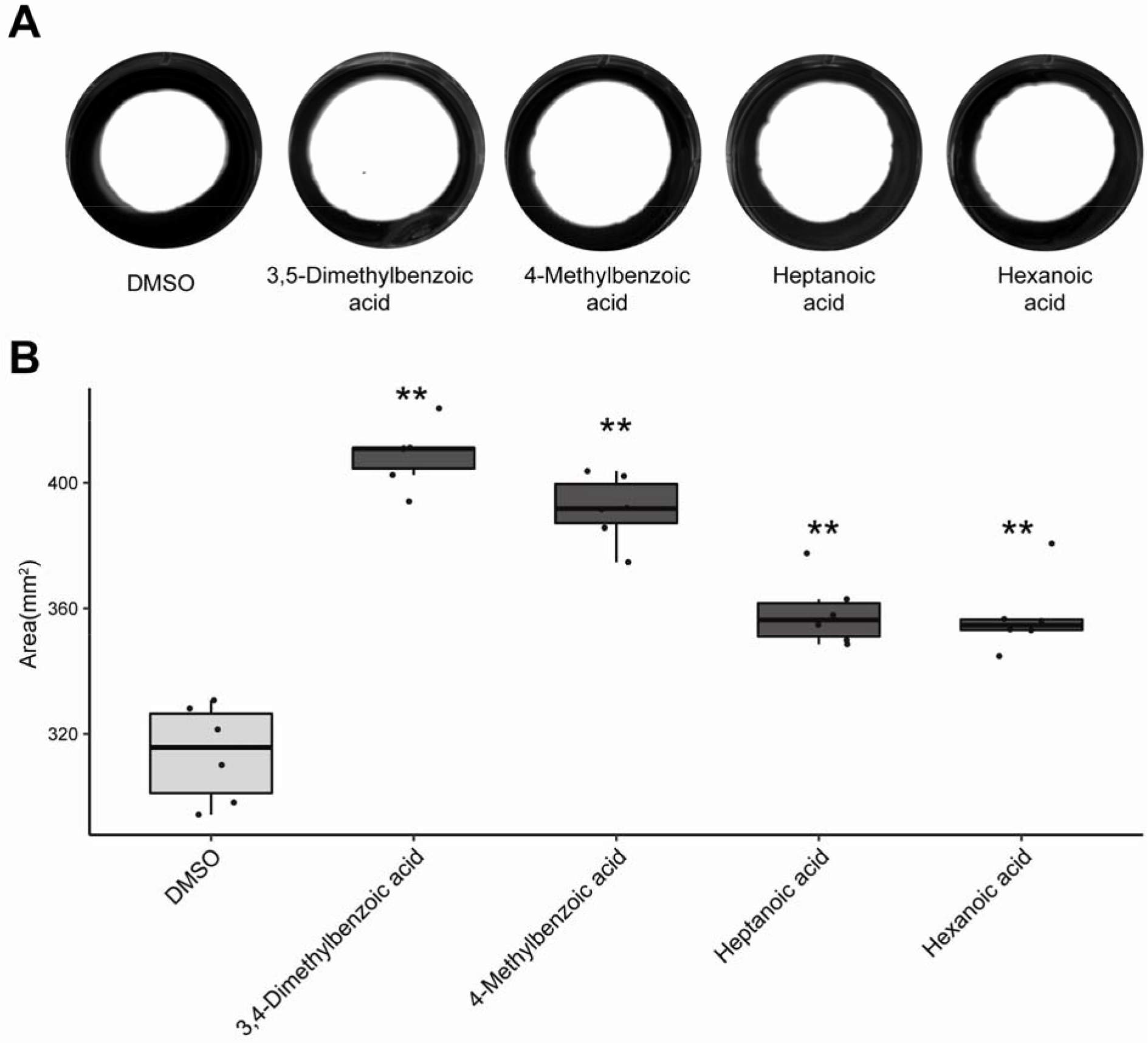
The 4 identified active metabolites increase EHEC motility in a plate-based swimming assay. (**A,B**). Effects of 3,4-dimethylbenzoic acid, 4-methylbenzoic acid, hexanoic acid, and heptanoic acid (all at 200 μM) individually on EHEC-GFP swimming motility. (**A**) Photographic image of the plate containing EHEC-GFP bacteria (white) cultured with each of the 4 metabolites. (B) Quantification of the area occupied by EHEC-GFP in **A**. **p<0.01.

## SUPPLEMENTARY TABLE LEGEND

**Additional Table 1.** List of 30 known metabolites enriched in Hmm compared to Mmm that were selected for *fliC*-luciferase screening (CAS n: Chemical Abstracts Service number; “name”: metabolites with a known name; “similarity”: closest MSMS spectrum in the reference database to the analyte, with a 95% confidence in identification).

**Additional Table 2.** Reagents and resources.

## REFERENCES

1. Cheng Y-L, Song L-Q, Huang Y-M, Xiong Y-W, Zhang X-A, Sun H, et al. Effect of enterohaemorrhagic Escherichia coli O157:H7-specific enterohaemolysin on interleukin-1β production differs between human and mouse macrophages due to the different sensitivity of NLRP3 activation. Immunology. 2015;145:258–67. doi:10.1111/imm.12442.

2. Perlman RL. Mouse models of human disease: An evolutionary perspective. Evol Med public Heal. 2016;2016:170–6. doi:10.1093/emph/eow014.

3. Chung H, Pamp SJSJ, Hill JA, Surana NK, Edelman SM, Troy EB, et al. Gut immune maturation depends on colonization with a host-specific microbiota. Cell. 2012;149:1578–93. doi:10.1016/j.cell.2012.04.037.

4. Eppinger M, Cebula TA. Future perspectives, applications and challenges of genomic epidemiology studies for food-borne pathogens: A case study of Enterohemorrhagic Escherichia coli (EHEC) of the O157:H7 serotype. Gut Microbes. 2015;6:194–201.

5. Mohawk KL, O’Brien AD. Mouse models of Escherichia coli O157:H7 infection and shiga toxin injection. J Biomed Biotechnol. 2011;2011:258185. doi:10.1155/2011/258185.

6. Keepers TR, Psotka MA, Gross LK, Obrig TG. A murine model of HUS: Shiga toxin with lipopolysaccharide mimics the renal damage and physiologic response of human disease. J Am Soc Nephrol. 2006;17:3404–14. doi:10.1681/ASN.2006050419.

7. Ikeda M, Ito S, Honda M. Hemolytic uremic syndrome induced by lipopolysaccharide and Shiga-like toxin. Pediatr Nephrol. 2004;19:485–9. doi:10.1007/s00467-003-1395-7.

8. Schüller S. Shiga toxin interaction with human intestinal epithelium. Toxins (Basel). 2011;3:626–39. doi:10.3390/toxins3060626.

9. Schüller S, Frankel G, Phillips AD. Interaction of Shiga toxin from Escherichia coli with human intestinal epithelial cell lines and explants: Stx2 induces epithelial damage in organ culture. Cell Microbiol. 2004;6:289–301. doi:10.1111/j.1462-5822.2004.00370.x.

10. Miyamoto Y, Iimura M, Kaper JB, Torres AG, Kagnoff MF. Role of Shiga toxin versus H7 flagellin in enterohaemorrhagic Escherichia coli signalling of human colon epithelium in vivo. Cell Microbiol. 2006;8:869–79. doi:10.1111/j.1462-5822.2005.00673.x.

11. Ritchie JM. Animal Models of Enterohemorrhagic Escherichia coli Infection. Microbiol Spectr. 2014;2:1–13. doi:10.1128/microbiolspec.EHEC-0022-2013.Correspondence.

12. Fukuda S, Toh H, Hase K, Oshima K, Nakanishi Y, Yoshimura K, et al. Bifidobacteria can protect from enteropathogenic infection through production of acetate. Nature. 2011;469:543–7. doi:10.1038/nature09646.

13. Jacobson A, Lam L, Rajendram M, Tamburini F, Honeycutt J, Pham T, et al. A Gut Commensal-Produced Metabolite Mediates Colonization Resistance to Salmonella Infection. Cell Host Microbe. 2018;24:296–307.e7. doi:10.1016/j.chom.2018.07.002.

14. Nakanishi N, Tashiro K, Kuhara S, Hayashi T, Sugimoto N, Tobe T. Regulation of virulence by butyrate sensing in enterohaemorrhagic Escherichia coli. Microbiology. 2009;155 Pt 2:521–30. doi:10.1099/mic.0.023499-0.

15. Tobe T, Nakanishi N, Sugimoto N. Activation of motility by sensing short-chain fatty acids via two steps in a flagellar gene regulatory cascade in enterohemorrhagic Escherichia coli. Infect Immun. 2011;79:1016–24. doi:10.1128/IAI.00927-10.

16. Surana NK, Kasper DL. Moving beyond microbiome-wide associations to causal microbe identification. Nature. 2017;552:244–7.

17. Sridharan G V., Choi K, Klemashevich C, Wu C, Prabakaran D, Pan L Bin, et al. Prediction and quantification of bioactive microbiota metabolites in the mouse gut. Nat Commun. 2014;5:1–13. doi:10.1038/ncomms6492.

18. Ingber DE. Reverse Engineering Human Pathophysiology with Organs-on-Chips. Cell. 2016;164:1105–9. doi:10.1016/j.cell.2016.02.049.

19. Jalili-Firoozinezhad S, Prantil-Baun R, Jiang A, Potla R, Mammoto T, Weaver JC, et al. Modeling radiation injury-induced cell death and countermeasure drug responses in a human Gut-on-a-Chip article. Cell Death Dis. 2018;9. doi:10.1038/s41419-018-0304-8.

20. Huh D, Leslie DC, Matthews BD, Fraser JP, Jurek S, Hamilton GA, et al. A human disease model of drug toxicity-induced pulmonary edema in a lung-on-a-chip microdevice. Sci Transl Med. 2012;4:159ra147. doi:10.1126/scitranslmed.3004249.

21. Benam KH, Novak R, Nawroth J, Hirano-Kobayashi M, Ferrante TC, Choe Y, et al. Matched-Comparative Modeling of Normal and Diseased Human Airway Responses Using a Microengineered Breathing Lung Chip. Cell Syst. 2016;3:456–466.e4. doi:10.1016/j.cels.2016.10.003.

22. Barrile R, van der Meer AD, Park H, Fraser JP, Simic D, Teng F, et al. Organ-on-Chip Recapitulates Thrombosis Induced by an anti-CD154 Monoclonal Antibody: Translational Potential of Advanced Microengineered Systems. Clin Pharmacol Ther. 2018. doi:10.1002/cpt.1054.

23. Kim HJ, Li H, Collins JJ, Ingber DE. Contributions of microbiome and mechanical deformation to intestinal bacterial overgrowth and inflammation in a human gut-on-a-chip. Proc Natl Acad Sci U S A. 2016;113:E7–15. doi:10.1073/pnas.1522193112.

24. Wang G, McCain ML, Yang L, He A, Pasqualini FS, Agarwal A, et al. Modeling the mitochondrial cardiomyopathy of Barth syndrome with induced pluripotent stem cell and heart-on-chip technologies. Nat Med. 2014;20:616–23. doi:10.1038/nm.3545.

25. Dawson A, Dyer C, Macfie J, Davies J, Karsai L, Greenman J, et al. A microfluidic chip based model for the study of full thickness human intestinal tissue using dual flow. Biomicrofluidics. 2016;10:064101. doi:10.1063/1.4964813.

26. Kasendra M, Tovaglieri A, Sontheimer-Phelps A, Jalili-Firoozinezhad S, Bein A, Chalkiadaki A, et al. Development of a primary human Small Intestine-on-a-Chip using biopsy-derived organoids. Sci Rep. 2018;8:2871. doi:10.1038/s41598-018-21201-7.

27. Zihler Berner A, Fuentes S, Dostal A, Payne AN, Vazquez Gutierrez P, Chassard C, et al. Novel Polyfermentor intestinal model (PolyFermS) for controlled ecological studies: validation and effect of pH. PLoS One. 2013;8:e77772. doi:10.1371/journal.pone.0077772.

28. Tanner SA, Zihler Berner A, Rigozzi E, Grattepanche F, Chassard C, Lacroix C. In vitro continuous fermentation model (PolyFermS) of the swine proximal colon for simultaneous testing on the same gut microbiota. PLoS One. 2014;9:e94123. doi:10.1371/journal.pone.0094123.

29. Fehlbaum S, Chassard C, Haug MC, Fourmestraux C, Derrien M, Lacroix C. Design and Investigation of PolyFermS In Vitro Continuous Fermentation Models Inoculated with Immobilized Fecal Microbiota Mimicking the Elderly Colon. PLoS One. 2015;10:e0142793. doi:10.1371/journal.pone.0142793.

30. Poeker SA, Geirnaert A, Berchtold L, Greppi A, Krych L, Steinert RE, et al. Understanding the prebiotic potential of different dietary fibers using an in vitro continuous adult fermentation model (PolyFermS). Sci Rep. 2018;8:4318. doi:10.1038/s41598-018-22438-y.

31. Andrews C, McLean MH, Durum SK. Cytokine Tuning of Intestinal Epithelial Function. Front Immunol. 2018;9:1270. doi:10.3389/fimmu.2018.01270.

32. Kagnoff MF. The intestinal epithelium is an integral component of a communications network. J Clin Invest. 2014;124:2841–3. doi:10.1172/JCI75225.

33. Maaser C, Heidemann J, von Eiff C, Lugering A, Spahn TW, Binion DG, et al. Human intestinal microvascular endothelial cells express Toll-like receptor 5: a binding partner for bacterial flagellin. J Immunol. 2004;172:5056–62. doi:10.4049/jimmunol.172.8.5056.

34. Eckmann L, Kagnoff MF, Fierer J. Epithelial cells secrete the chemokine interleukin-8 in response to bacterial entry. Infect Immun. 1993;61:4569–74. doi:<p></p>.

35. Shimizu M, Kuroda M, Sakashita N, Konishi M, Kaneda H, Igarashi N, et al. Cytokine profiles of patients with enterohemorrhagic Escherichia coli O111-induced hemolytic-uremic syndrome. Cytokine. 2012;60:694–700. doi:10.1016/j.cyto.2012.07.038.

36. Murata A, Shimazu T, Yamamoto T, Taenaka N, Nagayama K ichi, Honda T, et al. Profiles of circulating inflammatory- and anti-inflammatory cytokines in patients with hemolytic uremic syndrome due to E. coli O157 infection. Cytokine. 1998;10:544–8.

37. Fitzpatrick MM, Shah V, Trompeter RS, Dillon MJ, Barratt TM. Interleukin-8 and polymorphoneutrophil leucocyte activation in hemolytic uremic syndrome of childhood. Kidney Int. 1992;42:951–6. doi:10.1038/ki.1992.372.

38. López EL, Contrini MM, Devoto S, de Rosa MF, Graña MG, Genero MH, et al. Tumor necrosis factor concentrations in hemolytic uremic syndrome patients and children with bloody diarrhea in Argentina. Pediatr Infect Dis J. 1995;14:594–8. http://www.ncbi.nlm.nih.gov/pubmed/7567288.

39. van de Kar NC, Monnens LA, Karmali MA, van Hinsbergh VW. Tumor necrosis factor and interleukin-1 induce expression of the verocytotoxin receptor globotriaosylceramide on human endothelial cells: implications for the pathogenesis of the hemolytic uremic syndrome. Blood. 1992;80:2755–64. http://www.ncbi.nlm.nih.gov/pubmed/1333300.

40. Ceponis PJM, McKay DM, Ching JCY, Pereira P, Sherman PM. Enterohemorrhagic Escherichia coli O157:H7 disrupts Stat1-mediated gamma interferon signal transduction in epithelial cells. Infect Immun. 2003;71:1396–404. doi:10.1128/IAI.71.3.1396.

41. Proulx F, Toledano B, Phan V, Clermont M-J, Mariscalco MM, Seidman EG. Circulating granulocyte colony-stimulating factor, C-X-C, and C-C chemokines in children with Escherichia coli O157:H7 associated hemolytic uremic syndrome. Pediatr Res. 2002;52:928–34. doi:10.1203/00006450-200212000-00019.

42. Franke A, Balschun T, Karlsen TH, Sventoraityte J, Nikolaus S, Mayr G, et al. Sequence variants in IL10, ARPC2 and multiple other loci contribute to ulcerative colitis susceptibility. Nat Genet. 2008;40:1319–23. doi:10.1038/ng.221.

43. Marlow GJ, van Gent D, Ferguson LR. Why interleukin-10 supplementation does not work in Crohn’s disease patients. World J Gastroenterol. 2013;19:3931–41.

44. Fioranelli M, Roccia MG. Twenty-five years of studies and trials for the therapeutic application of IL-10 immunomodulating properties. From high doses administration to low dose medicine new paradigm. J Integr Cardiol. 2014;1:2—6.

45. Yamamoto T, Nagayama K, Satomura K, Honda T, Okada S. Increased serum IL-10 and endothelin levels in hemolytic uremic syndrome caused by Escherichia coli O157. Nephron. 2000;84:326–32. doi:10.1159/000045607.

46. Sassone-Corsi M, Raffatellu M. No vacancy: how beneficial microbes cooperate with immunity to provide colonization resistance to pathogens. J Immunol. 2015;194:4081–7. doi:10.4049/jimmunol.1403169.

47. Wadhams GH, Armitage JP. Making sense of it all: Bacterial chemotaxis. Nat Rev Mol Cell Biol. 2004;5:1024–37.

48. Lopes JG, Sourjik V. Chemotaxis of Escherichia coli to major hormones and polyamines present in human gut. ISME J. 2018;96:1272–82. doi:10.1038/s41396-018-0227-5.

49. Matilla MA, Krell T. The effect of bacterial chemotaxis on host infection and pathogenicity. FEMS Microbiol Rev. 2018;42:40–67. doi:10.1093/femsre/fux052.

50. Erhardt M. Strategies to Block Bacterial Pathogenesis by Interference with Motility and Chemotaxis. In: Stadler M, Dersch P, editors. How to Overcome the Antibiotic CrisisL: Facts, Challenges, Technologies and Future Perspectives. Cham: Springer International Publishing; 2016. p. 185–205. doi:10.1007/82_2016_493.

51. Ravichandar JD, Bower AG, Julius AA, Collins CH. Transcriptional control of motility enables directional movement of Escherichia coli in a signal gradient. Sci Rep. 2017;7:1–14. doi:10.1038/s41598-017-08870-6.

52. Lane MC, Alteri CJ, Smith SN, Mobley HLT. Expression of flagella is coincident with uropathogenic Escherichia coli ascension to the upper urinary tract. Proc Natl Acad Sci. 2007;104:16669–74. doi:10.1073/pnas.0607898104.

53. Valeriani C, Li M, Novosel J, Arlt J, Marenduzzo D. Colloids in a bacterial bath: simulations and experiments. Soft Matter. 2011;7:5228. doi:10.1039/c1sm05260h.

54. Hidalgo G, Chan M, Tufenkji N. Inhibition of Escherichia coli CFT073 flic expression and motility by cranberry materials. Appl Environ Microbiol. 2011;77:6852–7.

55. Tuttle J, Gomez T, Doyle MP, Wells JG, Zhao T, Tauxe R V, et al. Lessons from a large outbreak of Escherichia coli O157:H7 infections: insights into the infectious dose and method of widespread contamination of hamburger patties. Epidemiol Infect. 1999;122:185–92. doi:10.1017/S0950268898001976.

56. Gonthier M-P, Remesy C, Scalbert A, Cheynier V, Souquet J-M, Poutanen K, et al. Microbial metabolism of caffeic acid and its esters chlorogenic and caftaric acids by human faecal microbiota in vitro. Biomed Pharmacother. 2006;60:536–40. doi:10.1016/j.biopha.2006.07.084.

57. Andreasen MF, Christensen LP, Meyer AS, Hansen Å. Content of phenolic acids and ferulic acid dehydrodimers in 17 rye (Secale cereale L.) varieties. J Agric Food Chem. 2000;48:2837–42.

58. Ozdal T, Sela DA, Xiao J, Boyacioglu D, Chen F, Capanoglu E. The reciprocal interactions between polyphenols and gut microbiota and effects on bioaccessibility. Nutrients. 2016;8:1–36.

59. Lacroix C, Wouters T De, Chassard C. ScienceDirect Integrated multi-scale strategies to investigate nutritional compounds and their effect on the gut microbiota. Curr Opin Biotechnol. 2015;32:149–55. doi:10.1016/j.copbio.2014.12.009.

60. Zheng X, Qiu Y, Zhong W, Baxter S, Su M, Li Q, et al. A targeted metabolomic protocol for short-chain fatty acids and branched-chain amino acids. Metabolomics. 2013;9:818–27. doi:10.1007/s11306-013-0500-6.

61. Han J, Lin K, Sequeira C, Borchers CH. An isotope-labeled chemical derivatization method for the quantitation of short-chain fatty acids in human feces by liquid chromatography-tandem mass spectrometry. Anal Chim Acta. 2015;854:86–94. doi:10.1016/j.aca.2014.11.015.

62. Di Cagno R, De Angelis M, De Pasquale I, Ndagijimana M, Vernocchi P, Ricciuti P, et al. Duodenal and faecal microbiota of celiac children: molecular, phenotype and metabolome characterization. BMC Microbiol. 2011;11:219. doi:10.1186/1471-2180-11-219.

63. Bhatia SN, Ingber DE. Microfluidic organs-on-chips. Nat Biotechnol. 2014;32:760–72. doi:10.1038/nbt.2989.

64. Jalili-Firoozinezhad S, Gazzaniga FS, Calamari EL, Camacho DM, Fadel CW, Nestor B, et al. Complex human gut microbiome cultured in anaerobic human intestine chips. bioRxiv. 2018. http://biorxiv.org/content/early/2018/09/20/421404.abstract.

65. Vandussen KL, Marinshaw JM, Shaikh N, Miyoshi H, Moon C, Tarr PI, et al. Development of an enhanced human gastrointestinal epithelial culture system to facilitate patient-based assays. 2014.

66. Sato T, Clevers H. Growing self-organizing mini-guts from a single intestinal stem cell: mechanism and applications. Science. 2013;340:1190–4. doi:10.1126/science.1234852.

67. Schindelin J, Arganda-Carreras I, Frise E, Kaynig V, Longair M, Pietzsch T, et al. Fiji: an open-source platform for biological-image analysis. Nat Methods. 2012;9:676–82. doi:10.1038/nmeth.2019.

68. Tinevez J-Y, Perry N, Schindelin J, Hoopes GM, Reynolds GD, Laplantine E, et al. TrackMate: An open and extensible platform for single-particle tracking. Methods. 2017;115:80–90. doi:10.1016/j.ymeth.2016.09.016.

69. Thévenaz P, Ruttimann UE, Unser M. A pyramid approach to subpixel registration based on intensity. IEEE Trans Image Process. 1998;7:27–41. doi:10.1109/83.650848.

70. Yamaguchi N, Nasu M. Flow cytometric analysis of bacterial respiratory and enzymatic activity in the natural aquatic environment. 1997;:43–52.

71. Allison SE, Silphaduang U, Mascarenhas M, Konczy P, Quan Q, Karmali M, et al. Novel repressor of Escherichia coli O157:H7 motility encoded in the putative fimbrial cluster OI-1. J Bacteriol. 2012;194:5343–52.

72. R Team. R: A language and environment for statistical computing. R Foundation for Statistical Computing. 2016.

73. Wilcoxon F. Individual comparisons of grouped data by ranking methods. J Econ Entomol. 1946;39:269. doi:10.1093/jee/39.2.269.

74. Bonferroni CE. Teoria statistica delle classi e calcolo delle probabilità. Pubbl del R Ist Super di Sci Econ e Commer di Firenze. 1936.

75. Mann HB, Whitney DR. On a Test of Whether one of Two Random Variables is Stochastically Larger than the Other. Ann Math Stat. 1947;18:50–60. doi:10.1214/aoms/1177730491.

76. Bowne JB DLA. Metabolomics: Analysis of Metabolomics Data. 2014. https://cran.r-project.org/package=metabolomics.

77. Ritchie ME, Phipson B, Wu D, Hu Y, Law CW, Shi W, et al. limma powers differential expression analyses for RNA-sequencing and microarray studies. Nucleic Acids Res. 2015;43:e47. doi:10.1093/nar/gkv007.

78. Patro R, Duggal G, Love MI, Irizarry RA, Kingsford C. Salmon provides fast and bias-aware quantification of transcript expression. Nat Methods. 2017;14:417–9. doi:10.1038/nmeth.4197.

79. Dobin A, Davis CA, Schlesinger F, Drenkow J, Zaleski C, Jha S, et al. STAR: ultrafast universal RNA-seq aligner. Bioinformatics. 2013;29:15–21. doi:10.1093/bioinformatics/bts635.

80. Love MI, Huber W, Anders S. Moderated estimation of fold change and dispersion for RNA-seq data with DESeq2. Genome Biol. 2014;15:550. doi:10.1186/s13059-014-0550-8.

81. Yu G, Wang L-G, Han Y, He Q-Y. clusterProfiler: an R package for comparing biological themes among gene clusters. OMICS. 2012;16:284–7. doi:10.1089/omi.2011.0118.

